# An agent-built, verified atlas of public spatial transcriptomics shows that breadth across studies drives transfer and reproducibility

**DOI:** 10.64898/2026.05.27.727615

**Authors:** Jia-Hao Ji, Qi Zou, Jiabei Cheng, Zixi She, Yining Hao, Weishi Liu, Daoliang Zhang, Zhikang Wang, Jin-Tai Yu, Zhiyuan Yuan

**Affiliations:** Department of Neurology and National Center for Neurological Diseases, Huashan Hospital, State Key Laboratory of Medical Neurobiology and Ministry of Education Frontiers Center for Brain Science, Shanghai Medical College, Fudan University, Shanghai, 200040, China; Institute of Science and Technology for Brain-Inspired Intelligence, MOE Key Laboratory of Computational Neuroscience and Brain-Inspired Intelligence, MOE Frontiers Center for Brain Science, Fudan University, Shanghai, 200433, China; Center of Intelligent Medicine, School of Control Science and Engineering, Shandong University, Jinan, Shandong, China; School of Automation and Intelligent Sensing, Shanghai Jiao Tong University, Shanghai, 200433, China

## Abstract

Public spatial transcriptomics data are deposited study by study, in formats that differ from one laboratory to the next, and no resource has yet indexed or verified them as a whole. Here we present SpatialDataAgent-DB, an atlas built from the Gene Expression Omnibus (GEO) by SpatialDataAgent, a workflow in which language-model agents do the work and every output is checked rather than trusted. An assay index covering all 298,179 GEO series identifies 2,411 spatial series, which gives public spatial data an explicit denominator. For each series, an agent writes conversion code instead of editing the data directly, and a frozen, rule-based scorer recomputes every released value from the source files. In a held-out test, the scorer rejected all 1,231 corrupted copies. We also visually reviewed every released sample, including the 4,386 H&E-registered Visium samples of our earlier HESRT release. SpatialDataAgent-DB contains 12,163 samples from 12 platforms and 42 species in a single format, each with its provenance and defect flags. Its 8,211 image-paired samples are 5.9 times as many as in the next image-paired resource, and at least 71% of all samples are absent from five existing catalogues. Without batch correction, sequencing-based samples group by organ more than by platform: with platforms balanced, organ explains 32% of the variance of human pseudobulk profiles and platform 13% (mouse, 47% and 18%). We then used the atlas in two applications. Image-to-expression models trained on it transfer to unseen studies, and their accuracy rises steadily with training data and, at a fixed number of patches, with the number of training studies over the range tested (for H&E, Pearson *r* gains 0.0056 per doubling of training patches up to 524,288); 11 of 13 forecasts locked in advance were confirmed. Adding studies helps more than adding patches from the same studies, because each study leaves a fingerprint in its images. Intent-based search over the same index assembled 70 single-cell dementia studies, and the replication of disease signatures in held-out studies rose steadily with the number of studies (direction agreement averaged over 28 held-out studies, 63% with one study and 77% with 16). In both applications, performance kept rising with the number of independent studies, which a complete and verified atlas is built to provide.

## Introduction

Spatial transcriptomics measures gene expression in place within tissue, and within a decade it has advanced from 100 µm spots to subcellular imaging [1–8]. Most of these data are deposited in the Gene Expression Omnibus (GEO) [9], but GEO stores only what each laboratory chooses to upload. As a result, count matrices, coordinates, scale factors and images arrive in dozens of different layouts. Coordinate units are often unstated, and series descriptions mix the assays that were deposited with methods used elsewhere in the paper. A researcher who wants every public study that pairs spatial expression with an H&E image of the same section must therefore search by keyword and rebuild each dataset manually.

Automated resources have repeatedly changed what can be learned from public data. Examples include ARCHS4 and recount3 for bulk RNA-seq [10, 11], ChIP-Atlas for chromatin [12], and scBaseCount, in which language-model agents curate and reprocess single-cell data at scale [13]. Spatial databases, by contrast, remain curated collections of hundreds to a few thousand samples [14–17]. Machine-learning corpora such as HEST-1k and STimage-1K4M collect only what one model family needs [18, 19], as do collections built for spatial foundation models [20]. Repository metadata have long been mined to find samples [21], and language models now help to curate them [22]. Yet none of these resources indexes all of GEO, and none verifies each sample against its source files.

Spatial data are harder to process automatically than sequencing reads, because the spatial information sits in supplementary files: spot positions, their units, the alignment to an image, and the image itself. An agent that converts these files can fail silently. For example, it can flip an axis, apply the wrong scale or attach the matrix of another sample, and still produce a plausible dataset. Language-model agents can now plan and run scientific work [23–28], yet their output can be fluent but wrong [29]. The central design question is therefore not only whether an agent can perform the conversion, but how each output can be checked independently.

Here we build SpatialDataAgent-DB with the SpatialDataAgent workflow and put it to use. An index first labels every GEO series by assay, and a standardization agent then writes the mapping code for each spatial series. A frozen scorer, a fixed rule-based program rather than a language model, reexecutes this code against read-only source files. Through a second route, we add 4,386 H&E-registered Visium samples from our earlier HESRT release [30, 31]. Every released sample from both routes, 12,163 in total, is also reviewed visually. We then use the atlas as training data for models that predict gene expression from tissue images, measure how their accuracy scales with data and with the number of studies, and ask why these models transfer poorly to new studies. Finally, we test whether intent-based search over the index can assemble cohorts for a disease analysis. All three analyses point to one conclusion: independent studies provide breadth, which only a public atlas can supply.

Our contributions are:

- **A whole-GEO assay index.** We labelled all 298,179 GEO series by assay. The index makes the denominator of public spatial data explicit (2,411 series), so that recall and precision can be measured (Figures 1 and 2).
- **A verify-not-trust agent design.** The agent writes code against read-only sources, and an independent frozen scorer recomputes every released value. Across 12,326 conversions without human intervention, 91% passed after the agent repaired its own errors, and the validation gate rejected every corrupted test copy (Figure 2).
- **The largest verified image-paired spatial resource.** The resource contains 12,163 samples, 8,211 of them with a registered image. Most samples are absent from existing catalogues, and each carries its provenance and defect flags (Figure 3).
- **Evidence of what breadth is worth.** We ran a forecast-and-test training study on blind studies, which showed steady gains with data and with studies, and analysed why transfer to new studies fails (Figures 4 and 5). We also show an intent-search application that turns a single request into a 70-study dementia meta-analysis, whose replication also rises steadily with the number of studies (Figure 6).

**Figure 1.**
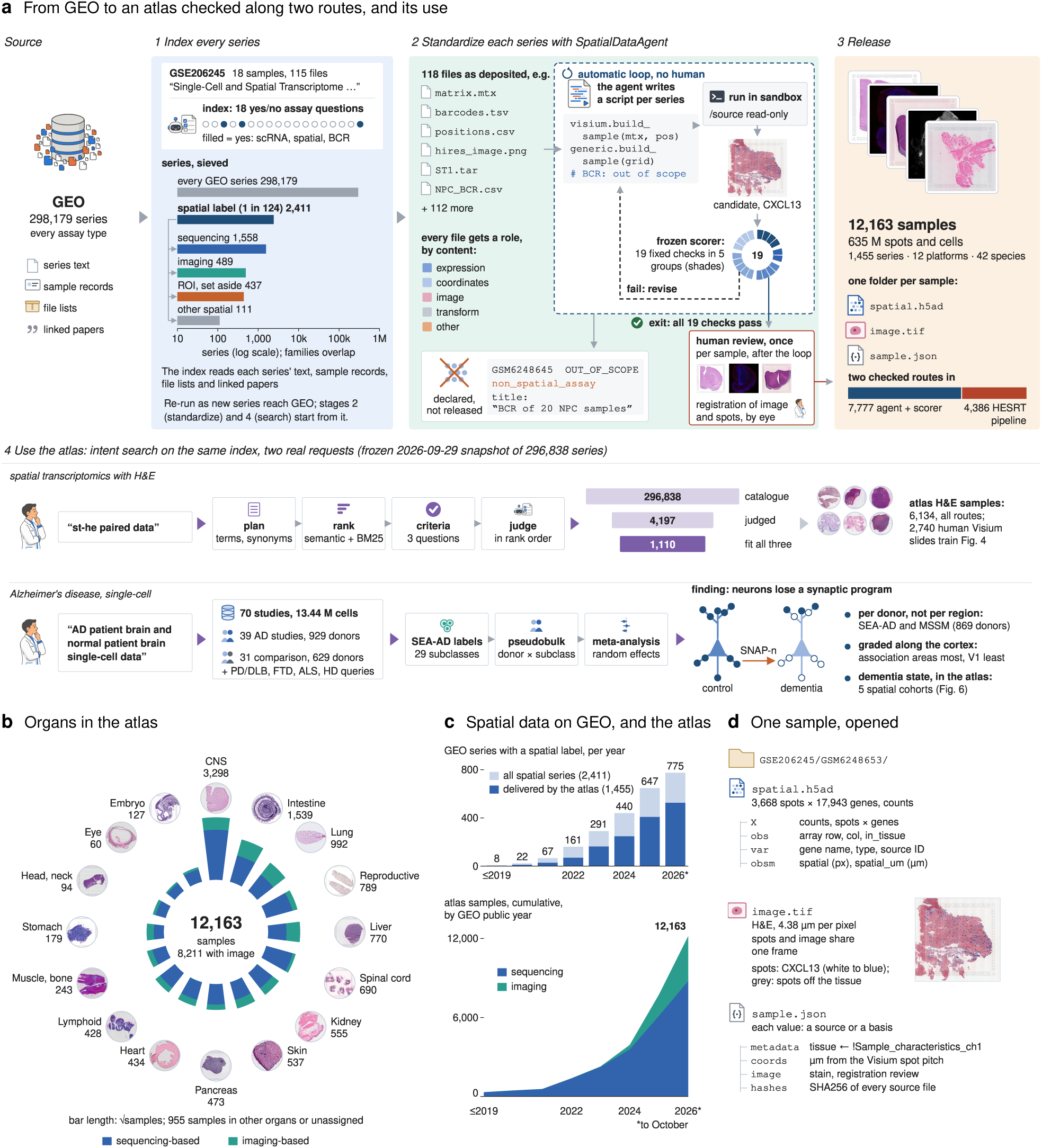
An agent-built, independently checked atlas of public spatial transcriptomics. (A) The workflow on one real series, GSE206245, and two real applications. (1) Index: every GEO series (298,179 in total) is read with its text, sample records, file lists and linked papers. A language model answers 18 yes/no assay questions per series, and a label is assigned when the probability is at least 0.5 (0.8 for imaging-based spatial assays). Bars filter the series on a log axis. In total, 2,411 series carry a spatial label, in overlapping families: sequencing (1,558), imaging (489), region of interest (437) and other spatial omics (111). (2) Standardize: in an automated loop without human input, the agent writes one script per series, which calls only shared readers and runs with the sources mounted read-only. A frozen scorer re-reads the sources and applies 19 fixed checks. A sample leaves the loop only when all checks pass or when it is marked unreleased; image registrations are then visually reviewed once. (3) Release: 12,163 samples from two checked routes, 7,777 delivered by the agent and scorer and 4,386 by the HESRT pipeline. (4) Use: intent-based search on the same index (“st-he paired data”; 1,110 matching series) and an application to single-cell data in Alzheimer’s disease (70 studies; analysis in Figure 6). (B) Organs in the atlas; bar length shows the square root of the sample count, split into sequencing- and imaging-based assays. (C) Top: GEO series with a spatial label by public release year (2026 up to 9 October); the dark portion marks series that deliver at least one standardized sample (1,455 of 2,411). Bottom: cumulative atlas samples. (D) One released sample, opened (GSE206245/GSM6248653). Shown are counts and positions in image pixels and micrometres, the H&E image in the same frame, and the record that cites a source or stated basis for every value.

**Figure 2.**
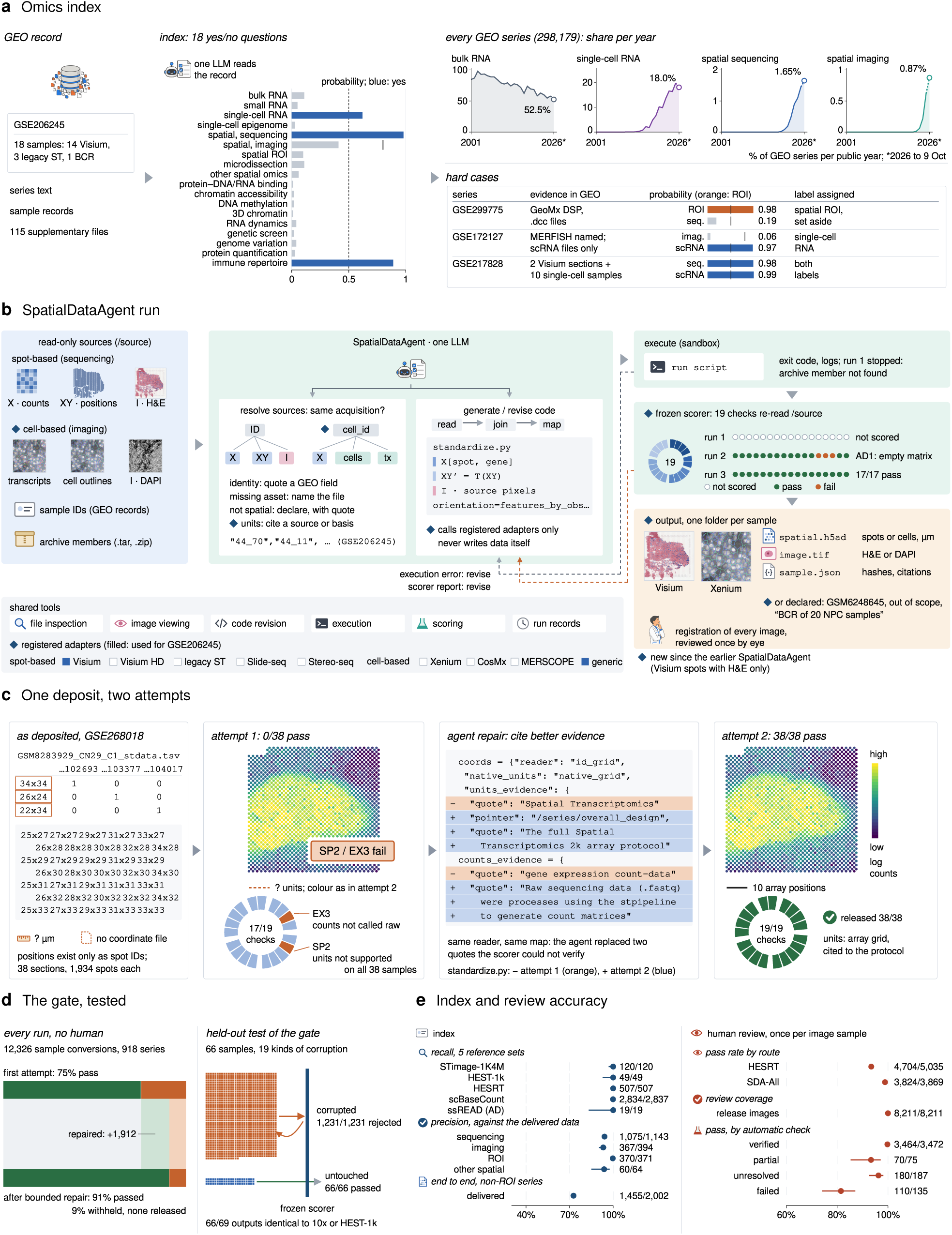
How the atlas is built: the index finds spatial series, and agent code is accepted only after recomputation. (A) The index on GSE206245: 18 answer probabilities, with thresholds as dashed lines (0.5 overall; 0.8 for imaging). Also shown are the share of GEO series carrying four labels in each year (2001–2026) and three hard cases. (B) A SpatialDataAgent run. Data sources are mounted read-only, and the agent resolves sample identities and units with cited evidence. The agent writes one script that calls only registered adapters; the frozen scorer then applies 19 checks in five groups and yields the delivered folders (Visium and Xenium examples). (C) One deposit over two attempts (GSE268018, legacy ST). The first attempt fails two citation checks (SP2, EX3) on all 38 samples; after the agent replaces two quotes, the second attempt passes 19 of 19 checks. (D) Left: all runs without human input, from the first scored attempt (9,299 of 12,326 conversions pass) to the final attempt. In total, 11,211 conversions pass; conversions are counted per run, so reruns of a series count again. The 1,115 withheld conversions were not released. The green band shows conversions recovered by bounded repair. Right: test of the gate on 66 held-out samples with 19 corruption types (one square per copy); the gate rejected all 1,231 corrupted copies and passed all 66 untouched copies. (E) Accuracy of the index and of review (dots with Wilson 95% CI). Recall on five reference sets (HEST-1k: 49 of 65 series; all 16 missed series are non-spatial companion datasets from the same papers). Precision against delivered data, for series where standardization reached a verdict. End to end, 1,455 of 2,002 non-region-of-interest spatial series delivered samples. Also shown are the human review pass rate by route, the coverage of released images, and the pass rate by status of the automatic registration check.

**Figure 3.**
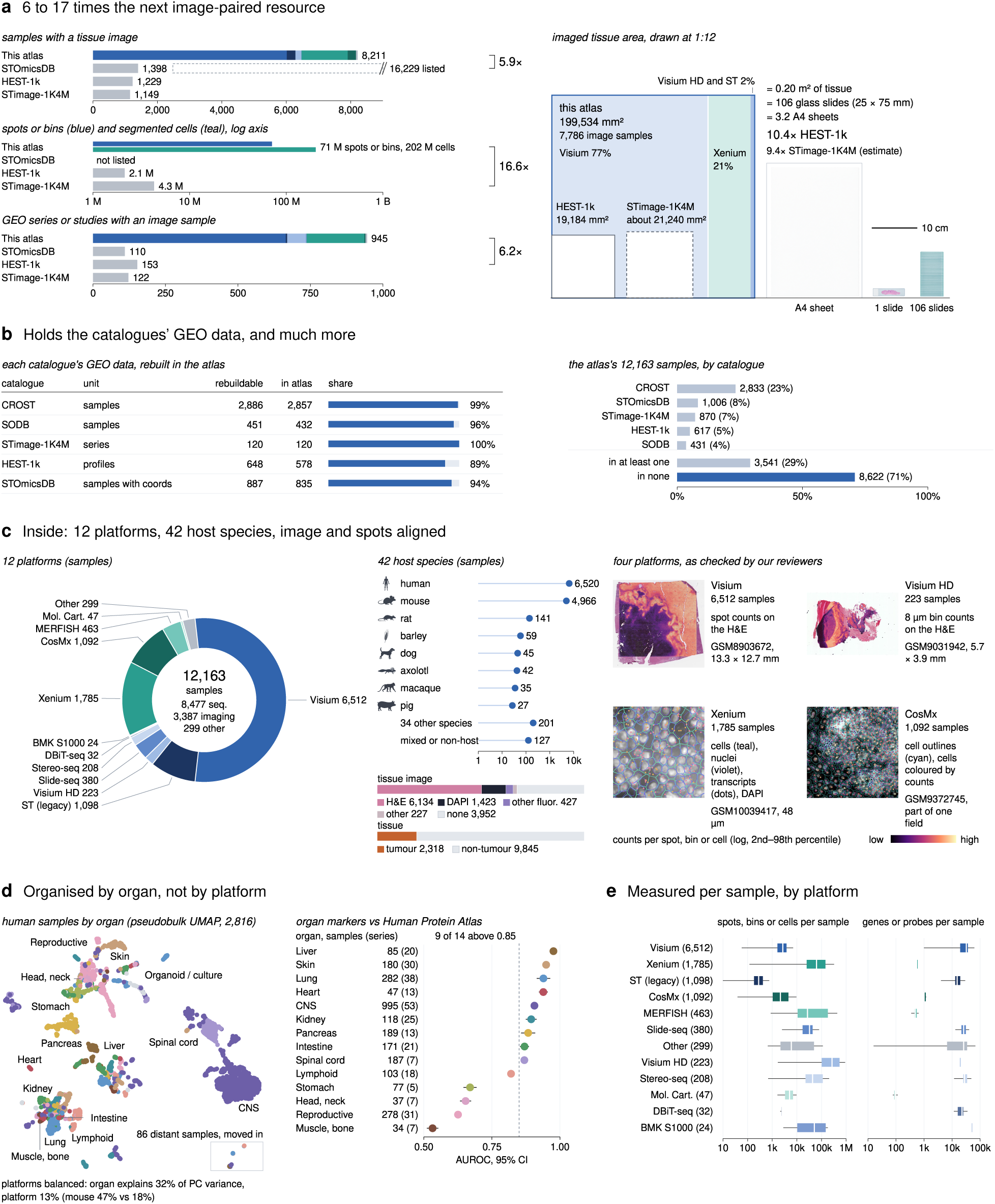
The atlas: 6 to 17 times the next image-paired resource, holding the catalogues’ data and much more, organised by organ. (A) Image-paired public resources compared with one counting rule per measure (September 2026). For samples with a tissue image, the dashed outline marks the 16,229 records listed by STOmicsDB, most without an image or coordinates. Also compared are spots or bins and segmented cells on image samples (log axis), series with an image sample, and all imaged tissue drawn at 1:12 (199,534 mm^2^, a lower bound from occupied 100 µm squares). (B) Left: each catalogue’s GEO data that can be rebuilt from GEO, and the share present in the atlas. Right: the 12,163 atlas samples grouped by catalogue membership; 8,622 samples (71%) are in none of the five catalogues (a lower bound, because HEST-1k and STimage-1K4M are matched by series). (C) Twelve platforms, 42 host species, image types and tumour status, with four platforms as checked by reviewers (Xenium: cells teal, nuclei violet, transcripts as dots). (D) Pseudobulk UMAP of 2,816 human sequencing-based samples without batch correction, coloured by organ. With platforms balanced, organ explains 32% of the variance of 50 principal components and platform 13% (mouse 47% and 18%; Figure S8A). Right: organ markers against the Human Protein Atlas (AUROC, 95% CI). (E) Spots, bins or cells and genes per sample, by platform.

**Figure 4.**
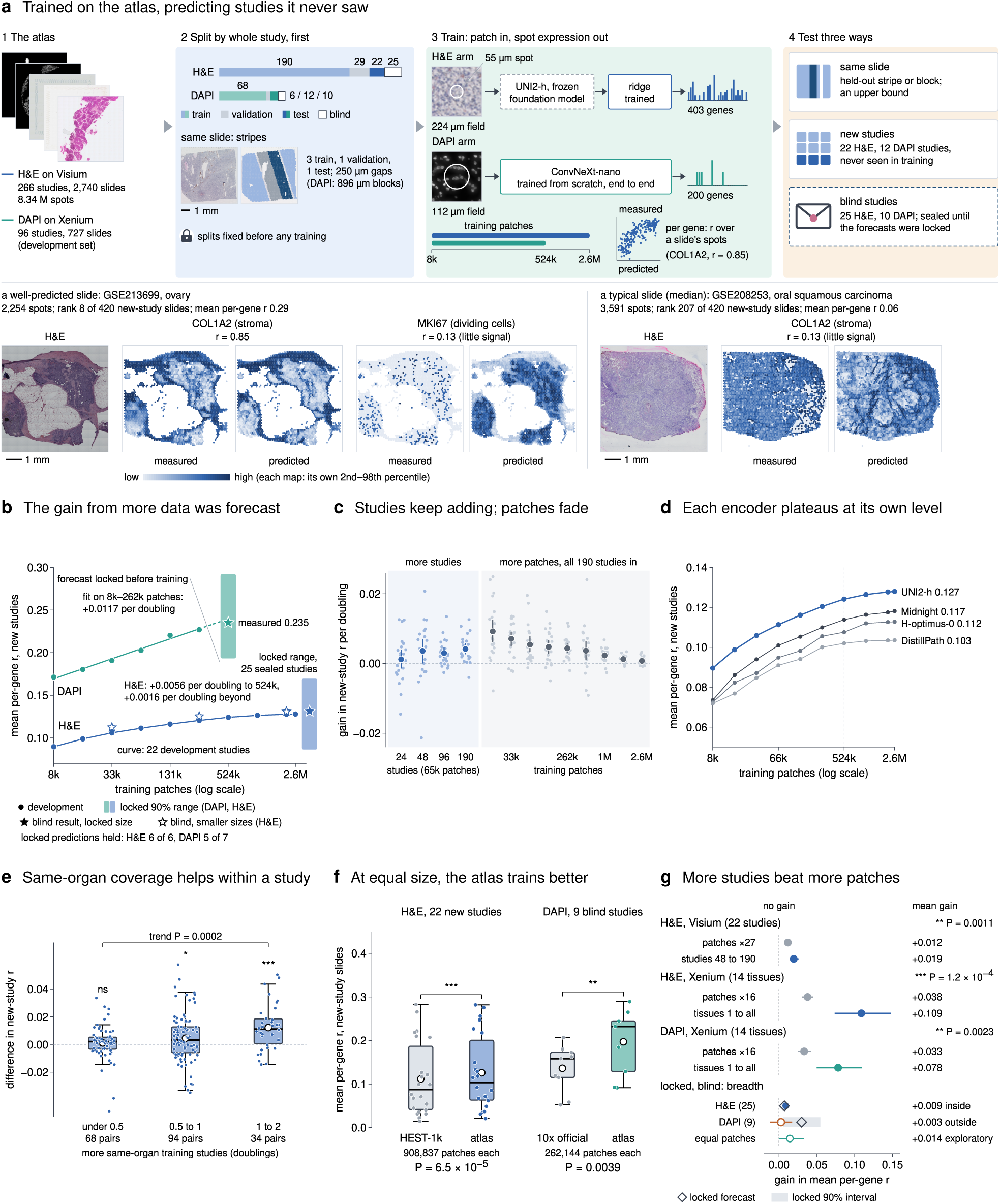
Trained on the atlas, image models predict expression in studies they never saw; accuracy rises with data, as forecast, and keeps rising with more studies. (A) Design. The atlas was split by whole study before training: the H&E arm used 190 training, 29 validation, 22 new-study test and 25 blind studies, and the DAPI arm 68, 6, 12 and 10. Shown are same-slide stripes, the two arms (H&E: frozen UNI2-h with a ridge read-out, 403 genes; DAPI: ConvNeXt-nano from scratch, 200 genes) and three evaluation tests. Bottom: measured and predicted expression on two new-study slides, a well-predicted slide (rank 8 of 420; mean per-gene *r* 0.29) and a typical slide (rank 207; 0.06, the median). (B) New-study *r* against training patches (log scale). DAPI: log-linear fit on 8k–262k patches, its extension to 524,288 patches (dashed), the locked 90% forecast range (band) and the result measured later (star). H&E: 22 development studies (dots), the locked range for the 25 blind studies, and blind results at the locked size (filled star) and at smaller sizes (open stars). Figure S4 shows all 13 locked predictions. (C) Gain in new-study *r* per doubling, for each successive doubling (faint dots, the 22 new studies; bold dot and line, mean and 95% bootstrap CI over studies). Left, more training studies at a fixed 65,536 patches (12 to 190 studies, mean of three random study orders). Labels mark the end of each doubling; for example, 24 marks the step from 12 to 24 studies. Right, more training patches in the registered order; all 190 studies are present from 32,768 patches, so later doublings add spots of the same slides (median 4 per slide at 8,192 patches, 277 at 524,288 and 1,374 at 2.6 million). (D) New-study *r* of four frozen pathology encoders (the fifth, Phikon-v2, is in Figure S9A), each with the same ridge read-out, training rows and test studies; the dashed line marks 524,288 patches. Labels give the mean over 1 to 2.6 million patches. (E) Same-organ coverage, within one test study. For each test study and number of training studies (12 to 96), two of the three random study orders are compared: the order with more training studies of the test study’s organ minus the one with fewer, binned by the difference in coverage (doublings of 1 + same-organ studies). Box plots show the pairs (IQR, median line, mean as open circle; dots, pairs). Marks above each box test the per-test-study mean difference against 0 (two-sided Wilcoxon signed-rank; ns, *P* = 0.55; *, *P* = 0.022; ***, *P* = 0.0006); the bracket gives the within-study trend. Across the 22 test studies, same-organ coverage did not predict new-study *r* (Spearman *ρ* = *−*0.27, *P* = 0.23). Within a test study, each doubling of same-organ training studies added 0.0067 (95% CI 0.0039–0.0090; bootstrap *P* = 0.0002; one fixed effect per test study and number of studies). (F) At equal size, the atlas trains better (one point per study; paired Wilcoxon). (G) Gain in new-study (or new-tissue) *r* from more patches of the same studies, against more studies at a fixed number of patches per slide (dots, mean; lines, 95% bootstrap CI). Bottom: locked comparisons on blind studies (diamond, forecast; band, locked 90% interval) and the exploratory equal-patch DAPI follow-up (open dot). Grey, more patches; coloured, more studies or tissues; orange ring, outside the locked interval.

**Figure 5.**
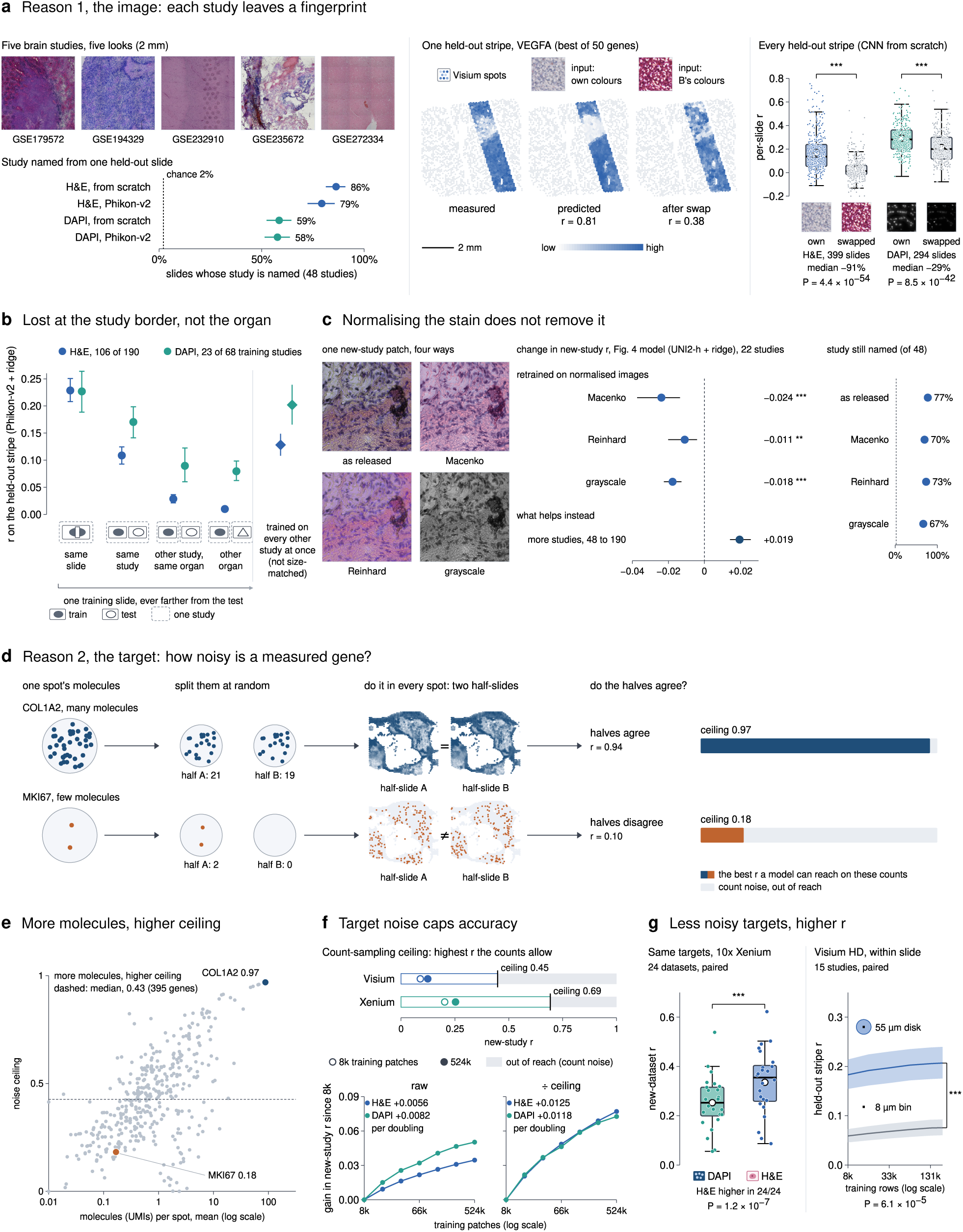
Two reasons transfer to new studies is hard: each study’s images carry a fingerprint that stain normalization does not remove, and Visium targets are noisy. (A) Brain sections from five studies. The study of a held-out slide is identified from image features alone (chance 2%). One held-out stripe is given the colour of another study (VEGFA, the best of the slide’s 50 most variable genes; *r* 0.81 to 0.38). For every held-out stripe, own against swapped colours (CNN from scratch; H&E 399 slides, DAPI 294). (B) Prediction from one training slide at increasing distance from the test slide (Phikon-v2 with a slide-centred ridge; 106 H&E and 23 DAPI studies with every rung available). Diamonds: training on every other study at once (not size-matched). (C) One patch under three normalizations. Change in new-study *r* when the model of Figure 4 is retrained on normalized slides (22 studies), compared with more studies, and the share of slides whose study is still identified. (D) Noise in the measured genes. The molecules of each spot are split at random into two in-silico half-samples; their agreement, corrected to full depth (Spearman–Brown), gives the ceiling, the highest *r* a model can reach against these counts. (E) Ceiling against molecules per spot for 395 genes on one slide. (F) Count-sampling ceiling over all genes (Visium 0.449, Xenium 0.692) and new-study *r* at 8k and 524k patches (frozen UNI2-h with ridge for both image types). Also shown is the gain in *r* since 8k patches, raw and divided by the ceiling. (G) Less noisy targets: H&E against DAPI on the same 10x Xenium targets (24 datasets), and one 8 µm bin against the 55 µm disk around it on Visium HD.

**Figure 6.**
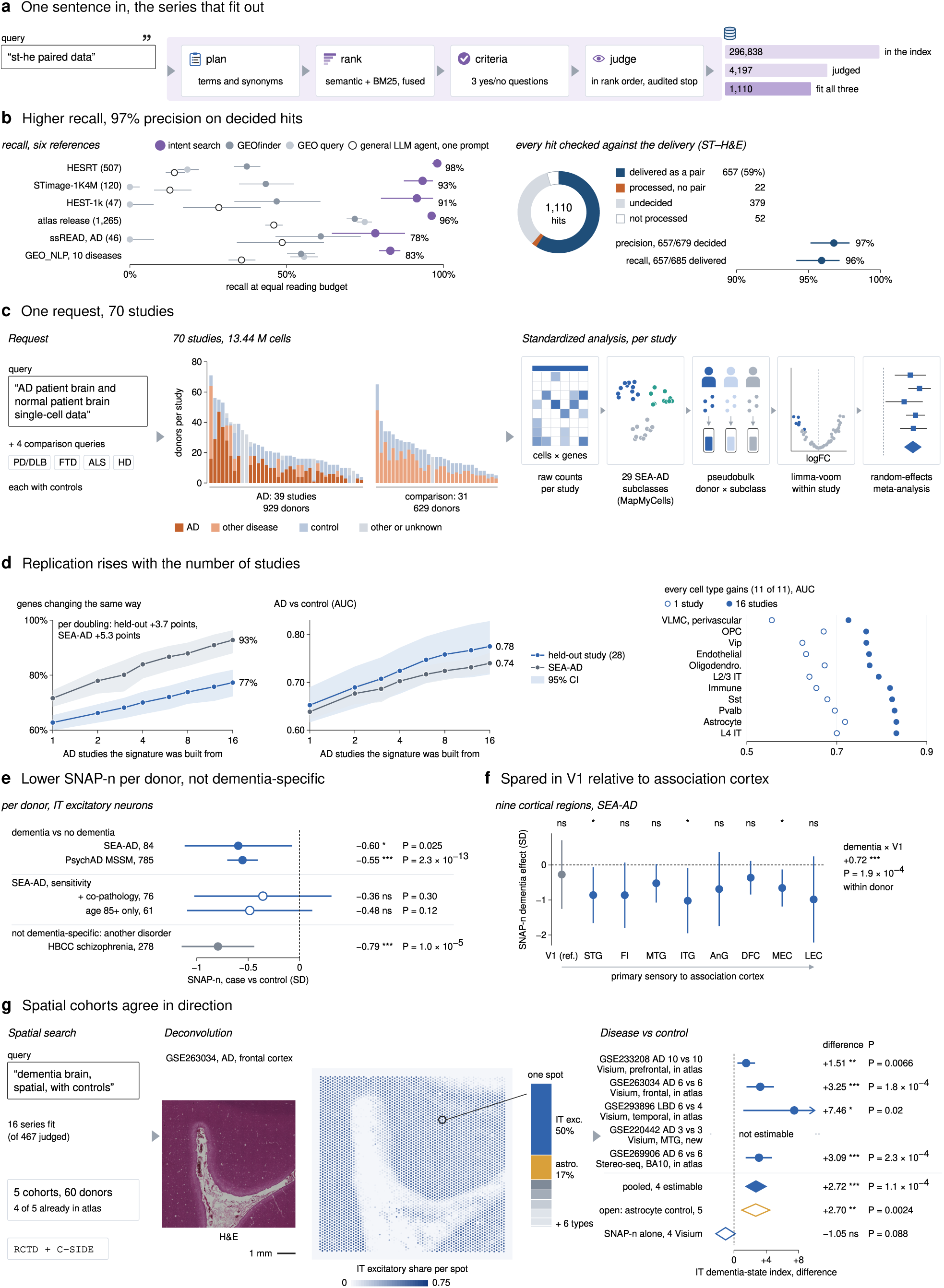
Intent search finds the data, and one request becomes a dementia analysis checked in the atlas. (A) Intent-based search for “st-he paired data” over 296,838 GEO series: plan, fused ranking, three generated criteria and rank-ordered judging (mechanics in Figure S6). (B) Left: recall on six independent references when every method reads as many series as intent search returns (Wilson 95% CI). Right: the 1,110 hits checked against the atlas’s deliveries (657 delivered as pairs, 22 without a pair, 379 undecided, 52 not processed); precision among hits with a verdict 97%, recall of delivered pairs 96%. (C) One request (and four comparison requests) yields 70 studies: donors per study by group, with every study processed by the same standardized pipeline. (D) Replication rises with the number of studies. A random-effects signature from *k* studies (*k* on a log scale) is tested on a study left out of the draw (each of 28 studies in turn) and on SEA-AD (bands, 95% CI); labels give the gain in direction agreement per doubling of *k*. Right: every cell type with 1 and 16 studies. (E) Donor-level SNAP-n in IT excitatory neurons, case against control (95% CI), including sensitivity analyses and schizophrenia. (F) SNAP-n dementia effect per region in SEA-AD; the decline is spared in V1 relative to association cortex (within-donor interaction). (G) A third search finds five spatial dementia cohorts. Shown are RCTD deconvolution of a released section and, per cohort, the difference in the IT dementia-state index with the pooled estimate, an astrocyte control (open amber diamond) and SNAP-n on its own (open blue diamond). GSE220442 (3 AD against 3 controls) gave no usable estimate of the IT state (standard error 183) and is shown but not pooled; the astrocyte control was estimable in all five.

## Results

### From GEO to an atlas checked along two routes

We built the resource in three stages (Figure 1A). First, an index covers every GEO series of every assay type, 298,179 series as of 9 October 2026. For each series, a language model reads the series text, sample records, file lists and linked papers, and answers 18 yes/no assay questions. GSE206245, for example, is labelled single-cell RNA, spatial sequencing and immune repertoire. In total, 2,411 series carry a spatial label, about 1 in 124. Their number has grown quickly, from 22 in 2020 to 647 in 2025 and 775 already in 2026 (Figure 1C). Second, a controller assigns a role to every deposited file based on its content. In an automated loop without human input, an agent writes one script per series; the script calls only shared readers and runs in a sandbox where sources are mounted read-only. A frozen scorer then re-reads the source files and applies 19 fixed checks. If any check fails, control returns to the agent. A sample leaves the loop only when it passes all 19 checks, or when it is declared unconvertible and not released. After the loop, we visually review the image–spot registrations. Third, we release 12,163 samples comprising 635 million spots and cells (Figure 1D). Each sample has its own folder. The folder holds a count matrix with coordinates (spatial.h5ad), together with a note, verified by the scorer, of whether the matrix contains raw counts. It also holds an image in the same coordinate frame (image.tif) and a record that cites a source or a stated basis for every value (sample.json). Each folder loads into AnnData and converts to SpatialData [32, 33]. A summary table of organ, species, platform, review status and defect flags lets users select samples before downloading.

The release comes from two verified routes: 7,777 samples passed the agent and scorer, and 4,386 H&E-registered Visium samples come from our earlier HESRT pipeline [30, 31]. We visually reviewed every sample from both routes. Of the 2,411 spatial series, 1,455 provide at least one sample. Of these, 1,265 contribute under their own GEO identifier and 190 through a linked SuperSeries, SubSeries or earlier deposit (Figure 1C). The same index also supports intent-based search for requests that predefined labels cannot express (Figure 1A, bottom; Figure 6).

### The index finds spatial series, and agent code is accepted only after recomputation

The index reports each answer as a probability and assigns a label when the probability is at least 0.5 (Figure 2A). For imaging-based spatial assays the threshold is 0.8, because papers often mention these assays without depositing their data. Across all of GEO, the labels show how deposits have changed over time. In 2026, bulk RNA accounts for 52.5% of new series and single-cell RNA for 18.0%, whereas spatial sequencing accounts for 1.65% and spatial imaging for 0.87%. The questions also help to resolve hard cases. A GeoMx deposit of .dcc files is labelled as region of interest, an assay type that falls outside the coordinate standard of the release. A series whose paper mentions MERFISH, but which deposits only single-cell matrices, is not labelled as imaging. A series with two Visium sections and ten single-cell samples receives both labels.

We evaluated the index in two ways (Figure 2E). Recall was tested on five external reference sets built without the index. The spatial labels recalled all 120 STimage-1K4M series, all 507 HESRT series, and 49 of the 65 GEO series linked to HEST-1k. All 16 missed series were non-spatial companion datasets from the same papers. The single-cell label recalled 2,834 of 2,837 human scBaseCount series and all 19 ssREAD series [34]. Precision was assessed against the final standardized outputs, counting a label as correct if the series yielded samples of that technology or, for region-of-interest labels, if the run confirmed such an assay. Among series that reached a verdict, the label was correct for 1,075 of 1,143 sequencing-labelled series (94%), 367 of 394 imaging-labelled series (93%) and 370 of 371 region-of-interest series. Overall, 1,455 of the 2,002 non-region-of-interest spatial series delivered standardized samples. Most of the remaining series lack coordinate or expression files on GEO (320) or have not yet been run (78).

The standardization agent does not create or modify data directly; instead, it resolves the data sources for each sample (Figure 2B). The agent determines which files belong to each sample, joins them by spot or cell identifier, and records physical units with a quoted source or a stated reason. It then writes one script per series, which calls only generic readers and registered adapters for five spot-based and three cell-based platforms. For example, on series GSE206245 the agent ran 56 shell commands. Its first run halted before scoring and the second passed 14 of 17 eligible samples; after a one-line fix, the third passed all 19 checks for all 17 samples. Code repairs often resolve missing evidence rather than numeric errors (Figure 2C). GSE268018, a legacy ST dataset, stores positions only as spot identifiers. The scorer rejected all 38 samples on the first attempt because two citations did not support the stated units or raw counts. After the agent replaced these quotes with sentences from the protocol and data-processing sections, the second attempt passed.

Across all runs without human intervention, the scorer evaluated 12,326 sample conversions from 918 series (Figure 2D). Of these, 75% passed on the first attempt and 91% after bounded automated repair, which recovered 1,912 conversions. The 1,115 conversions that still failed or produced no output were withheld. Conversions are counted per run, so a series rerun under a newer scorer version is counted again. The release keeps one passing conversion per sample, which gives 7,777 agent-route samples after deduplication. To test the gate itself, we corrupted 66 correctly standardized samples with 19 types of error, from shuffled spots and flipped axes to shifted gene labels and broken citations (Figure S3A). The frozen scorer rejected all 1,231 corrupted copies and passed all 66 untouched samples (Figure 2D; Figure S3B). Against external ground truth, 66 of 69 outputs matched the vendor 10x Genomics files or HEST-1k deposits exactly, and the other three differed only in which edge spots were retained (Figure S1D). Every released sample was also reviewed visually, both the 8,211 samples with a registered image and the 3,952 without one. Review decides whether an image is released. In the HESRT review cohort, 4,704 of 5,035 image samples passed, and 4,386 of the passing samples are released through the HESRT route. On the agent route, 3,824 of 3,869 reviewed image registrations passed and 4 were marked uncertain (Figure 2E). The 3,825 agent-route image samples in the release are the 3,824 that passed and one uncertain registration that the automatic check had verified. An image whose registration failed review is never released; the sample is either withheld or released with its expression and coordinates only.

### The largest public image-paired spatial resource, organised by organ

The atlas holds 8,211 samples with a registered tissue image, 5.9 times as many as the next image-paired resource. On every shared measure, the atlas is 6 to 17 times larger than its neighbours (Figure 3A). These image samples contain 71 million spots or bins and 202 million segmented cells from 945 series, against 153 series in the next resource. Imaged tissue covers 199,534 mm^2^, 10.4 times HEST-1k counted the same way, whereas most of the 16,229 records listed by STOmicsDB have no image or coordinates. The atlas also contains almost all of the catalogues’ data that can be rebuilt from GEO (Figure 3B). Coverage is 99% for CROST, 96% for SODB, 100% of STimage-1K4M series, 89% for HEST-1k (every series was found; the rest lie in three series whose files the pipeline cannot read yet) and 94% for STOmicsDB samples with coordinates. Conversely, 8,622 of the 12,163 released samples (71%) appear in none of the five catalogues. Because two catalogues can be matched only by series, this count is a lower bound.

The release spans 12 platforms and 42 host species (Figure 3C), at resolutions from 100 µm legacy spots to single cells [1, 4, 7, 8, 35, 36]. The release includes 6,134 H&E and 1,423 DAPI images and 2,318 tumour samples. Spatial structure is present on every platform: in 10,879 of 12,093 evaluable samples (90%), the median Moran’s I exceeded its maximum under coordinate permutation (Figure S9B). The micrometre scale was verified from the source files for 5,151 samples and taken from a platform constant for 4,236; for another 24, it was declared but not verified. The remaining 2,752 samples keep their native units (pixels, array grid or unstated), carry no micrometre coordinates and are flagged (Figure S9C). Recomputation also exposed defects in the deposits themselves (Figure S2D), which we recorded as per-sample flags rather than repairing them silently. For example, 363 samples state no coordinate unit, 105 of 1,785 Xenium samples carry a periodic field-of-view grid in their counts, and 77 samples were re-deposited in a later series. These defects matter for reuse: an unflagged re-deposit can leak between training and test sets, and a grid can masquerade as spatial structure.

Without batch correction, the standardized sequencing-based samples group more by organ than by platform (Figure 3D), although platform effects remain and are largest for CosMx (Figure S8C). Because Visium accounts for 92% of the 2,816 human sequencing-based samples (274 series), we compared the two factors on draws in which each organ contributes the same number of series per platform. In this balanced design, organ explained 32% of the variance of the first 50 principal components and platform 13% (Figure S8A), against 13% and 5% for shuffled labels. In mouse, organ explained 47% of the variance and platform 18%. Platform effects are therefore real, but organ effects are larger. Organ markers ranked across series agreed well with the Human Protein Atlas [37], with an AUROC above 0.85 for 9 of 14 organ systems. Within single platforms, organ labels predicted from samples of other studies exceeded the majority baseline for the sequencing platforms and for mouse Xenium. For human Xenium (69% against 54%) and CosMx (77% against 64%), accuracy was close to baseline, with confidence intervals that include the baseline, and few genes are shared across studies (Figure S8B). Organ labels also transferred from Visium to other platforms. The 15 nearest Visium samples from other series predicted the organ of 91% of mouse Slide-seq and 84% of mouse Stereo-seq samples. They also predicted the organ of 74% (human) to 85% (mouse) of Xenium samples, although mouse Xenium shared fewer than 90 genes with Visium. Transfer failed for CosMx (11% human, 7% mouse), whose profiles lay far from every Visium sample, and for human Slide-seq, whose samples are mostly brain (Figure S8C). These checks hold at the level of organs; they do not show that finer cell states can be compared without integration.

### Trained on the atlas, image models transfer to unseen studies, and accuracy rises with data, as forecast

We next asked whether the atlas is useful training data for models that predict spatial expression from tissue images [38–42] (Figure 4A). In the H&E arm (2,740 Visium slides from 266 studies), a frozen pathology foundation model, UNI2-h [43], encoded a 224 µm field around each 55 µm spot. A ridge read-out then predicted the expression of 403 genes. UNI2-h was the strongest of five frozen encoders, and larger MLP read-outs on its features did not exceed the ridge read-out (Figure S9A). In the DAPI arm (727 Xenium slides from 86 studies), a ConvNeXt-nano network [44] trained from scratch read a 112 µm field to predict 200 genes for a 55 µm pseudo-spot. The atlas was split by whole study before training, so every model was scored on studies it had never seen. A further 25 H&E and 10 DAPI studies were kept blind and sealed until we had committed forecasts of their results; one DAPI blind study was later removed by a grid filter before scoring, which left 9 evaluable DAPI studies. On a well-predicted slide from a new study (ovary, rank 8 of the 420 slides in the 22 new-study test studies), the model recovers the stromal gene COL1A2 (Pearson *r* = 0.85 across the spots of the slide) but not the proliferation marker MKI67 (*r* = 0.13). Averaged over all 403 genes, the median new-study slide reaches only *r* = 0.06; the mean over slides and studies reported below is higher because some slides are predicted well. Because the score is a correlation across the spots of one slide, any prediction that ignores the image, such as mean tissue expression, scores 0. Transfer is therefore real but modest, and it follows tissue structure: a gene’s new-study *r* tracks its spatial autocorrelation within a slide (Spearman *ρ* = 0.68; Figure S9E), so stromal and extracellular-matrix programmes transfer best and immune and cell-cycle genes least. Part of the remaining shortfall lies in the counts themselves (Figure 5F).

Over the range tested, new-study accuracy rose steadily with training data, and the gain was forecast before it was measured: 11 of 13 predictions locked before the blind studies were opened held (Figure 4B; Figure S4). For the DAPI network, new-study *r* rose by 0.0117 per doubling of training patches between 8,192 and 262,144. A log-linear curve fitted on these points predicted a score of 0.240 (90% interval 0.193–0.292) at 524,288 patches, and the model trained afterwards reached 0.235. For H&E, new-study *r* rose from 0.090 with 8,192 patches to 0.124 with 524,288, by 0.0056 per doubling on average (95% CI 0.0042–0.0070). Beyond this size the H&E curve flattened: each doubling added 0.0016 (0.0012–0.0021), and the model reached 0.128 with 2.6 million patches. Over these ranges, a log-linear curve, a power law and a power law in the error fit the DAPI, study and dementia curves about equally well, so we report average gains per doubling rather than a functional form. Only the full H&E patch curve separates the shapes: a saturating power law fitted on 8,192 to 262,144 patches predicted the four larger sizes, whereas a log-linear curve overshot them by 0.010 (Figure S4C). On the 25 blind studies, the 2.6-million-patch model scored 0.131, inside the locked range (0.086–0.169). The 13 locked predictions covered performance levels, gains, diminishing returns, encoders, study breadth, network size, and image against expression-only controls; all 6 held for H&E and 5 of 7 for DAPI.

Two limits fit this plateau, and these data cannot separate them: the fixed set of training studies and the frozen encoder. Two other explanations, the count ceiling and the study fingerprint, do not fit it (Figure 4C,D; Figure S9D). First, training patches were added in a fixed random order, so 189 of the 190 training studies were already present at 8,192 patches and all of them from 32,768. Each later doubling therefore added spots from the same slides, from a median of 277 per slide at 524,288 patches to 1,374 at 2.6 million, and the gain per doubling fell to 0.0007 (Figure 4C). In contrast, raising the number of training studies (and with them the number of slides) from 12 to 190 at a fixed 65,536 patches added 0.0030 per doubling (0.0017–0.0045), and the gain per doubling did not decline: the last doubling, from 96 to 190 studies, added 0.0041 (0.0026–0.0056). Second, four frozen encoders levelled off at the same data size but at different heights, and UNI2-h stayed 0.010–0.024 above the other three (higher in 20 of 22 new studies; Figure 4D). MLP read-outs on UNI2-h features stayed below the ridge read-out (Figure S9A). Because all four frozen encoders levelled off at the same data size, a limit of frozen features cannot be excluded, and the DAPI network, trained end to end rather than frozen, was still on its forecast trend at 524,288 patches. More studies at full patch depth, or an H&E encoder trained end to end, would be needed to separate the two limits. Third, the plateau lies far below the count-sampling ceiling of the Visium targets (0.449; defined in Figure 5D), and same-slide accuracy, measured on held-out stripes of training slides, levelled off as well, from 0.150 to 0.153 between 524,288 and 2.6 million patches. The ratio of new-study to same-slide accuracy stayed at 0.83–0.85 at every size (Figure S9D), so the plateau does not arise at the border between studies.

Which studies are added matters as well (Figure 4E). In the runs that varied the number of studies, the three random study orders gave each test study a different number of training studies from its own organ at the same total. Within a test study, each doubling of same-organ training studies added 0.0067 to its new-study *r* (95% CI 0.0039–0.0090; bootstrap *P* = 0.0002), about as much as doubling all training patches (0.0056), and the effect held without the seven brain test studies (+0.0051, 0.0009– 0.0085). Across test studies, by contrast, organ coverage did not predict accuracy (Spearman *ρ* = *−*0.27, *P* = 0.23). Brain, the best-covered organ with 30 training studies, was among the hardest to predict. Coverage of an organ therefore helps a given study, but it does not make every tissue equally predictable.

At equal size, the atlas was better training data than existing corpora (Figure 4F). We compared 908,837 patches from each source, with the same encoder, read-out, target genes and test studies. The atlas reached 0.126 against 0.112 for HEST-1k (+0.014, 95% CI 0.009–0.019; higher in 18 of 22 new studies; paired Wilcoxon *P* = 6.5 *×* 10*^−^*^5^). For DAPI at 262,144 patches, models trained on the atlas outperformed those trained on the official 10x Genomics Xenium data on all 9 evaluable blind studies, scoring 0.197 against 0.136 (+0.061, 0.032–0.095), in line with our prior forecast.

A controlled test confirmed that adding studies helped more than adding patches (Figure 4G). In the H&E arm, raising the number of studies from 48 to 190 at 64 patches per slide increased new-study *r* by 0.019 (0.014–0.026). Using all patches from the 48 studies instead gave 27 times as many patches, several times the total of the 190-study arm, yet raised new-study *r* by only 0.012 (0.009–0.015; paired *P* = 0.0011). A similar pattern held on 35 public Xenium sections with post-Xenium H&E from 14 tissue groups. At a fixed 8,192 patches, training on all other tissues rather than one gained 0.109 for H&E and 0.078 for DAPI. In contrast, 16 times more patches of the same tissue gained only 0.038 (H&E; paired *P* = 1.2 *×* 10*^−^*^4^) and 0.033 (DAPI; *P* = 0.0023). On the blind studies, the H&E breadth gain (+0.009) fell inside its locked interval. The locked DAPI breadth prediction, however, failed: the advantage of all studies was 0.003, below its locked interval (0.005–0.054). Because this design gave the arm with fewer studies three to five times more patches in total, it confounded breadth with data size. In an exploratory follow-up with equal numbers of patches in both arms, more studies helped by +0.036 on the development studies (12 of 12). On the blind studies, the gain was +0.014 (90% CI 0.002–0.030; 7 of 9 studies), although this result was not significant (*P* = 0.16).

### Two reasons transfer is hard: study fingerprints and count noise

Transfer to new studies loses accuracy for two reasons. The first lies in the images (Figure 5A). From image features alone, a classifier identified which of 48 training studies a held-out slide came from. The classifier was correct in 86% of cases with an H&E network trained from scratch and in 79% with frozen Phikon-v2 features (DAPI, 59% and 58%; chance, 2%). These study signatures echo the site signatures known from computational pathology [45]. Giving a held-out stripe the colour histogram of another study collapsed prediction: across 399 H&E slides, the median *r* fell from 0.135 to 0.013 (*−*91%), compared with *−*29% for DAPI. Accuracy is lost at the border between studies rather than between organs (Figure 5B). A model trained on one slide predicted another stripe of the same slide with *r* = 0.229 and another slide of the same study with 0.109. Its accuracy fell to 0.029 for a slide from another study of the same organ and to 0.010 for another organ. By comparison, training on every other study at once, with far more data, reached 0.128.

Stain normalization, the standard remedy [46], does not remove the fingerprint (Figure 5C; Figure S5). We normalized every slide with Macenko [47] or Reinhard [48], or converted it to grayscale, and then re-encoded the slides and retrained the model of Figure 4. New-study *r* fell under all three normalizations (Macenko *−*0.024, higher in 0 of 22 studies, *P* = 4.8 *×* 10*^−^*^7^), and the classifier still identified the study from the features in 67–73% of slides. Because a frozen encoder might simply read normalized slides poorly, we also trained an encoder by self-distillation to ignore changes in colour, stain, blur and resolution. This encoder cut the colour-swap loss from 33% to 9% but did not change new-study *r* either. Of all the strategies we tested, only more training studies helped (+0.019), in line with the steady gain with the number of studies (Figure 4C).

The second reason lies in the target data (Figure 5D–G). Splitting the molecules of each spot at random into two halves in silico gives two half-samples of the same section. Their agreement, corrected to full depth [49, 50], sets a noise ceiling: the highest *r* that any model can reach against these counts, given count sampling. For COL1A2 the halves agree (*r* = 0.94; ceiling 0.97), whereas for MKI67 they do not (*r* = 0.10; ceiling 0.18). Across 395 genes, the ceiling rises with the number of molecules per spot (median 0.43). Over all genes, it is 0.449 for the Visium targets of the H&E arm and 0.692 for the Xenium targets of the DAPI arm (Figure 5F). With the same frozen UNI2-h encoder and ridge read-out on both image types, DAPI gained more per doubling of data in raw *r* (0.0082 against 0.0056 for H&E). Once each gain is divided by the noise ceiling of its arm’s targets, however, the two gains are comparable (H&E 0.0125, 95% CI 0.0095–0.0156; DAPI 0.0118, 0.0095–0.0142). On the same targets, H&E is the better image. On 24 public Xenium datasets with post-Xenium H&E, models trained on H&E predicted new datasets better than models trained on DAPI in all 24 (0.336 against 0.253; *P* = 1.2 *×* 10*^−^*^7^; Figure 5G).

### Intent search finds the data, and one request becomes a dementia analysis

Fixed assay labels cannot express a request such as spatial transcriptomics paired with an H&E image of the same section. Intent-based search turns such a sentence into search plans, ranks all GEO series by fusing semantic and BM25 rankings [51–55], writes its own yes/no criteria, and judges series in rank order. Judging continues to a minimum depth and then stops when new matches run out; an audit below the stopping point checks for misses (Figure 6A; Figure S6). For “st-he paired data”, the search judged 4,197 series and returned 1,110 that met all three criteria. For comparison, every method was given the same reading budget, defined as the number of series returned by intent search. Within this budget, intent search recalled 78–98% of six independent reference sets (Figure 6B). The semantic search engine GEOfinder recalled 37–72% [56], an expert GEO query 0–75% and general-purpose language-model agents 13–49%. Checked against the atlas’s own deliveries, 657 of the 679 hits with a verdict were delivered H&E pairs (97%), and the search found 657 of the 685 delivered pairs (96%). This completeness has a price: one search cost US$12.4 at the prices used, about 100 times a general-purpose agent query. Intent search is therefore suited to broad requests with many matching series rather than to narrow ones (Limitations).

Although single-cell studies of the Alzheimer’s disease (AD) brain each profile tens of donors [57–60], their findings agree only in part [61]. We gave the search a research request, “AD patient brain and normal patient brain single-cell data”, together with four comparison requests for other neurodegenerative diseases (Figure 6C). The search assembled 70 studies with 13.44 million cells: 39 AD studies (929 donors) and 31 comparison studies (629 donors). Every study was processed with the same pipeline, which mapped cells to the SEA-AD subclasses [62, 63], built donor-by-subclass pseudobulks [64], ran limma-voom within each study [65, 66], and combined the studies by random-effects meta-analysis [67].

Breadth again decided the result (Figure 6D). We built the meta-analysis from *k* randomly drawn studies and tested it on a study left out of the draw. Each of the 28 AD studies with at least four donors in both groups served in turn as the test study. The share of signature genes that changed in the same direction rose steadily with the number of studies, from 63% with one study to 77% with 16, by 3.7 percentage points per doubling on average (95% CI 3.0–4.3). The donor-level AUC for disease against control rose from 0.65 to 0.78 and increased in every one of 11 cell types. In the independent SEA-AD atlas, agreement rose from 72% to 93%, by 5.3 points per doubling (4.7–5.8).

The main component of the excitatory-neuron dementia programme was a decline of SNAP-n, a synaptic programme described by Ling et al. [68]. The down-regulated genes overlapped its top 200 genes (odds ratio 10–22; Figure S7C). Synapse loss is the strongest structural correlate of cognitive decline in AD [69]. Per donor, SNAP-n was lower with dementia in SEA-AD (*−*0.60 SD, *P* = 0.025, 84 donors) and in PsychAD MSSM (*−*0.55, *P* = 2.3 *×* 10*^−^*^13^, 785 donors) [70] (Figure 6E). The signal is not specific to dementia: as Ling et al. reported, it fell as much in schizophrenia (*−*0.79, *P* = 1.0 *×* 10*^−^*^5^). In SEA-AD, the decline was no longer significant after adjustment for other brain pathologies (*P* = 0.30). The decline varied mainly between donors, with 81% of the variance of donor-by-region scores lying between donors. Within a donor, SNAP-n did not track local pTau or A*β*, whereas an astrocyte control tracked local pTau (Figure S7A,B,D). Across nine cortical regions, SNAP-n was spared in primary visual cortex relative to association cortex (dementia *×* V1 interaction +0.72, *P* = 1.9 *×* 10*^−^*^4^; Figure 6F). Proteins encoded by SNAP-n genes also correlated with cognition in an independent proteome [71] (Figure S7G). Among the comparison diseases, Parkinson’s disease with dementia and dementia with Lewy bodies largely shared the AD signature, but its neuronal part appeared only with dementia (Figure S7I,J).

Finally, a third search found five spatial dementia cohorts with cases and controls [72], four of them already in the atlas (Figure 6G). We deconvolved the spots of each section to estimate expression within each cell type, rather than from cell-type proportions [73, 74]. The broader excitatory dementia-state index shifted in the same direction in every estimable cohort (pooled +2.72, 95% CI 1.35–4.10). An astrocyte state, however, shifted just as much, and SNAP-n on its own did not reach significance (*−*1.05, *−*2.25 to 0.15, *P* = 0.088). In spot-level data, the disease signal is therefore not specific to neurons. SNAP-n on its own could be tested only in the four Visium cohorts, and the Visium count ceiling (Figure 5F) offers one explanation for this result. A cell-resolved spatial cohort would provide a direct test.

## Discussion

We built a resource in which agents do the laborious work of finding and converting public spatial data, but none of their output is accepted without verification. The index makes the denominator explicit, the agent writes only code that a frozen scorer re-executes against the read-only sources, and image registrations are reviewed visually. Judging generated code by running it rather than by reading it is also how code-writing models are evaluated [75, 76]. This division of labour lets the agent work at the scale of GEO: across 12,326 conversions it repaired most of its own failures, and in a held-out test the gate rejected every corrupted copy. The result is the largest verified image-paired resource of public spatial data, with 5.9 times as many image-paired samples as the next resource and at least 71% of its samples absent from five existing catalogues. Recomputation also exposed defects that would otherwise reach users silently, such as periodic field-of-view grids in 105 Xenium samples and 77 re-deposited samples that could leak between training and test sets. The same design distinguishes the atlas from curated spatial databases [14–17] and from machine-learning corpora [18, 19]: it has an explicit denominator, verifies every sample, and is re-run as GEO grows. Whereas scBaseCount showed the value of agents for single-cell reads [13], spatial data also require that positions, units and images be verified, not only recounted.

As training data, the atlas is both larger and more useful than existing corpora. At equal size, it trained better models than HEST-1k (higher in 18 of 22 new studies) and than the official 10x Xenium data (all 9 blind studies). New-study accuracy rose steadily with data, and 11 of 13 predictions locked before the blind studies were opened held, so these gains were forecast rather than fitted after the fact [77–79]. The training study also shows where the gains come from. At a fixed number of patches per slide, raising the number of studies from 48 to 190 gained more than using 27 times as many patches from the 48 studies. For a given test study, each doubling of training studies from its own organ added about as much as doubling all training patches, and once every study was present, more patches from the same slides added little. Pathology foundation models are now trained on ever larger slide collections [43, 80–82]. Our results suggest that, for transfer to new studies, the number and diversity of studies matter at least as much as the number of patches, in line with work on vision models in which data diversity sets robustness more than quantity [83, 84]. Because public data are the only practical source of hundreds of independent studies, an atlas of public data, and a search that can assemble cohorts from it, are worth building. Spatial deposits in GEO are growing quickly (647 series in 2025 and 775 by 9 October 2026), and the atlas is re-run as GEO grows; if the gain with studies holds beyond the range tested, each re-run should make the atlas more useful as training data. Several curve shapes fit these ranges about equally well (Figure S4C), so we report an empirical trend rather than a scaling law, and we report the locked prediction that failed alongside those that held.

The atlas also made the remaining gap measurable, and it has two sources. Every study leaves a fingerprint in its images that models can read, and transfer is lost at the border between studies. Stain normalization did not remove this fingerprint and lowered new-study accuracy in every case, so the fingerprint is not simply colour; it behaves like the batch effects long known in genomics [85, 86]. The counts themselves also cap the attainable correlation, at about 0.45 for Visium targets against 0.69 for Xenium targets. Once this ceiling is taken into account, H&E images carry as much learnable signal per doubling of data as DAPI images, and on the same Xenium targets H&E predicted new datasets better than DAPI in all 24. Image-to-expression models are therefore likely to gain from targets with less count noise as well as from more studies.

The dementia analysis shows what an agent-assisted study can reach. One request assembled 70 single-cell studies, and signatures built from more studies replicated better, both in held-out studies and in SEA-AD. The meta-analysis replicated a known synaptic programme, SNAP-n, which was lower with dementia in 785 PsychAD donors, and located its decline in the cortex: the decline was spared in primary visual cortex relative to association cortex. The same analysis also set limits on the finding. The programme fell as much in schizophrenia, so it is not specific to dementia, and spot-level spatial data could not resolve it in neurons; the Visium count ceiling offers one explanation, and a cell-resolved spatial cohort would provide a direct test.

## Limitations

Several limitations apply to the resource and to the analyses built on it. The atlas standardizes the processed files that authors deposited rather than re-quantifying raw reads, so differences between upstream pipelines remain. The micrometre basis cannot be verified for 2,752 samples, which keep their native units. Region-of-interest assays measure regions rather than spots or cells and fall outside the coordinate standard of the release. The 4,386 HESRT samples come from our earlier pipeline and were checked by human review, not by the agent’s scorer. Index precision was judged only on series whose standardization reached a verdict. The share of atlas samples absent from existing catalogues is a lower bound, and the comparison covers five catalogues. In the training study, the blind DAPI set contained 9 evaluable studies, and the locked DAPI breadth prediction failed; the equal-patch follow-up was run after this miss and is exploratory. These trends hold over the ranges tested, their functional form cannot be identified from these ranges (8,192 to 2.6 million patches and 12 to 190 training studies; 1 to 16 studies in the dementia meta-analysis), and in the training study more studies also meant more slides, so the two cannot be fully separated. The noise ceiling covers count sampling only. In the dementia analysis, most comparison datasets have no pathology staging for their donors, so we cannot separate a shared dementia process from AD co-pathology [87]. The spatial check of this analysis was also made on spot-level data. Finally, intent-based search costs more per request than a general-purpose agent (US$12.4 against under US$0.15), mostly because of a fallback model for very long series. In return, it found 96% of the confirmed series for a broad request, where agents found 3–14%. Intent search is therefore suited to broad requests with many matching datasets. A very specific request with several conditions can return no series at all (in a six-condition islet query, the best candidate met four). In that case, a cheaper agent query may serve as well.

## Methods

The methods are summarized at the level of design. Every run reported here is registered with its model identifiers, prompts and parameters. The prompts, the agent’s tool library, the scorer’s implementation and the analysis code are available to editors and reviewers on request, and they will be released with the peer-reviewed version.

### Whole-GEO assay index

For every GEO series (298,179 public up to 9 October 2026), we assembled the series text, sample records, supplementary file lists and linked papers. A language model answered 18 overlapping yes/no assay questions, each with a probability. The questions cover bulk, single-cell and spatial transcriptomics, spatial imaging, region-of-interest and other spatial omics, chromatin, methylation, screens, immune repertoire and others. A label is set at a probability of at least 0.5 (0.8 for imaging-based spatial assays), and the 2,411 series with any spatial label form the candidate pool. Recall was measured on five reference sets built without the index. Precision was judged against the final product among series whose standardization reached a verdict; a label counted as correct when the series delivered samples of that technology.

### Verify-not-trust standardization

Four design rules make agent output checkable. (1) *Read-only sources*: before the agent starts, the supplementary files of each series are frozen with content hashes and assigned roles based on their content. (2) *Code, not data*: the agent writes one script per series that may call only a closed library of platform readers and typed transforms. Numeric parameters must come from the readers, and every metadata value must cite a verbatim source passage or a stated basis. (3) *An independent frozen scorer*: the scorer was written and fixed before the runs, and it re-reads the source files without the agent’s code. It applies 19 checks in five groups (source and identity, matrix structure, expression values, coordinates, and provenance and citations), rebuilds the observations and compares every released value. The agent may repair its script for a bounded number of scored rounds, after which the controller reruns the scorer itself and releases only passing samples. (4) *Human review*: image–spot registrations are reviewed visually, and re-deposits and aliases are resolved in one final release-wide pass. We tested the gate by copying 66 correctly standardized samples once per corruption type (19 types) and scoring each copy once. We also compared its outputs with vendor and HEST-1k outputs for the same deposits.

### Atlas characterization

Catalogue coverage was matched by GEO sample, or by series for HEST-1k and STimage-1K4M, which gives a lower bound. For sequencing-based samples, pseudobulk profiles were reduced to 50 principal components without batch correction. The variance explained by organ or by platform is the *R*^2^ of one-hot labels. We computed this *R*^2^ on draws in which every organ contributes equal numbers of series per platform, and compared it with labels shuffled within the other factor. Organ prediction within and across platforms used genes measured in every compared sample, and organ markers were scored against the Human Protein Atlas [37].

### Forecast-and-test training study

Before any training, the atlas was split by whole study, and 25 H&E and 10 DAPI studies were kept blind until forecasts had been committed. H&E models used a frozen pathology encoder with a per-gene ridge read-out (403 genes); DAPI models were convolutional networks trained from scratch on Xenium pseudo-spots (200 genes). Performance was the Pearson *r* of each gene across the spots of a slide, averaged over genes, slides and studies. Forecasts for larger training sets and for the blind studies were fitted on development results only, locked, and scored once. Breadth and depth were varied at a fixed number of patches per slide, at equal totals, and on public Xenium sections with post-Xenium H&E. Scaling trends are the least-squares gain in new-study *r* per doubling of training size, with 95% intervals from bootstrapping test studies. Log-linear, power-law, power-law-in-the-error and saturating power-law curves were compared by AICc and by predicting the larger sizes from fits on the smaller ones. Organ coverage was the number of training studies of the test study’s organ (the most common organ of its slides); its within-study effect used the variation between the three random study orders at the same number of studies, with one fixed effect per test study and number of studies and 95% intervals from bootstrapping test studies. Patches were added in the registered order, and studies in three random orders at 65,536 patches.

### Study fingerprints and the count ceiling

Study identity was predicted directly from image features. Colour swaps, stain normalizations [47, 48] and an appearance-invariant encoder were tested by retraining the read-out. The count-sampling ceiling splits the molecules of each spot binomially into two halves and corrects their agreement to full depth with the Spearman–Brown formula [49, 50].

### Intent-based search

Given a request in plain language, the system converts it into search plans and fuses semantic and BM25 rankings of all GEO series [53, 55]. The system then writes its own yes/no criteria, and a judging model reads the series in rank order until new matches run out. In the benchmarks, every method read as many series as intent search returned.

### Dementia application

Every study returned by the search was processed with the same pipeline. Cells were mapped to SEA-AD subclasses [62, 63], donor-by-subclass pseudobulks were computed [64], and limma-voom was applied within each study [65, 66], followed by random-effects meta-analysis across studies [67]. To test reproducibility, we built a signature from *k* randomly drawn AD studies, using the top 50 up- and 50 down-regulated genes per cell type. Each signature was scored on a held-out study and on SEA-AD. We then measured the share of genes changing in the same direction and the AUC of a donor score for AD against control. Gains with *k* are the least-squares slope on log_2_ *k*, with 95% intervals from bootstrapping held-out studies (SEA-AD: cell types). SNAP-n [68] was scored per donor in IT excitatory neurons, adjusting for pathology, age and sex. Spatial cohorts were deconvolved with RCTD and C-SIDE [73, 74].

### Statistics

Proportions are reported with 95% Wilson intervals [88]. Paired comparisons used two-sided Wilcoxon signed-rank tests with study-bootstrap 95% intervals. Forecast intervals are 90%. All reported numbers were read directly from registered result files.

## Data and code availability

The release is available at https://huggingface.co/datasets/Biogod/SpatialDataAgent-DB. It contains 12,163 sample folders, sample and study tables, a schema and a verification report. Each sample can be read as AnnData and converted to SpatialData [32, 33]. The agent, scorer, prompts, index labels and analysis code will be released at https://github.com/Biogod2020/spatial-data-agent-db with the peer-reviewed version.

## Acknowledgments

Z.Y. acknowledges support from the National Natural Science Foundation of China (grant numbers 62303119 and 32470706), Shanghai Science and Technology Development Funds (grant number 23YF1403000), Fund of Fudan University and Cao’ejiang Basic Research (grant number 24FCA10), and the Computational Biology Program (number 25JS2850200) of the Science and Technology Commission of Shanghai Municipality (STCSM).

## Author contributions

J.-H.J. conceived the study, developed the SpatialDataAgent framework, performed the data mining, agentic curation, standardization, benchmarking, statistical analysis and visualization, and wrote the manuscript. Q.Z., J.C. and Z.S. reproduced the experiments and validated the computational results. Y.H., W.L. and D.Z. contributed to dataset checking, result organization and technical validation. Z.W. contributed to the conceptual framing of the agentic methodology, reviewed the computational workflow and revised the manuscript. J.-T.Y. provided biomedical supervision and computational resources. Z.Y. supervised the study, guided the overall manuscript structure, reviewed the computational framework and revised the manuscript.

---

**Figure S1.**
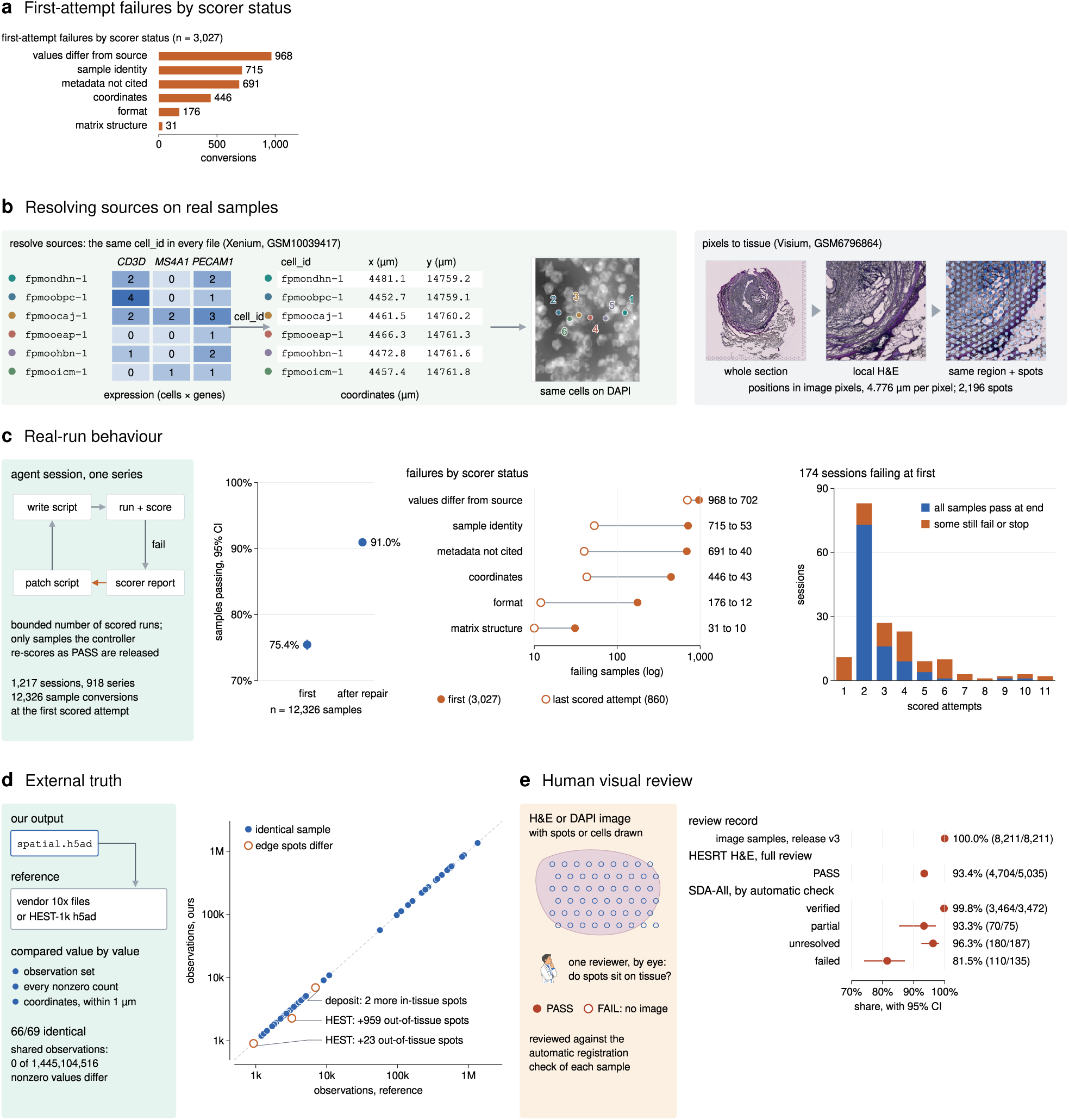
Agent runs and how their outputs are checked, related to Figure 2. (A) First-attempt failures by scorer status across 3,027 conversions. (B) Resolving data sources on real samples: a Xenium sample joined by cell identifier across expression, coordinates and the DAPI image, and a Visium sample with spots aligned on the H&E image. (C) Agent behaviour in real runs: success at the first and last attempt, failures by status, and sessions by number of scored attempts. (D) External ground truth: 66 of 69 outputs were identical to vendor or HEST-1k outputs. (E) Human visual review of image–spot registration; uncertain reviews count as not passed, as in Figure 2E.

**Figure S2.**
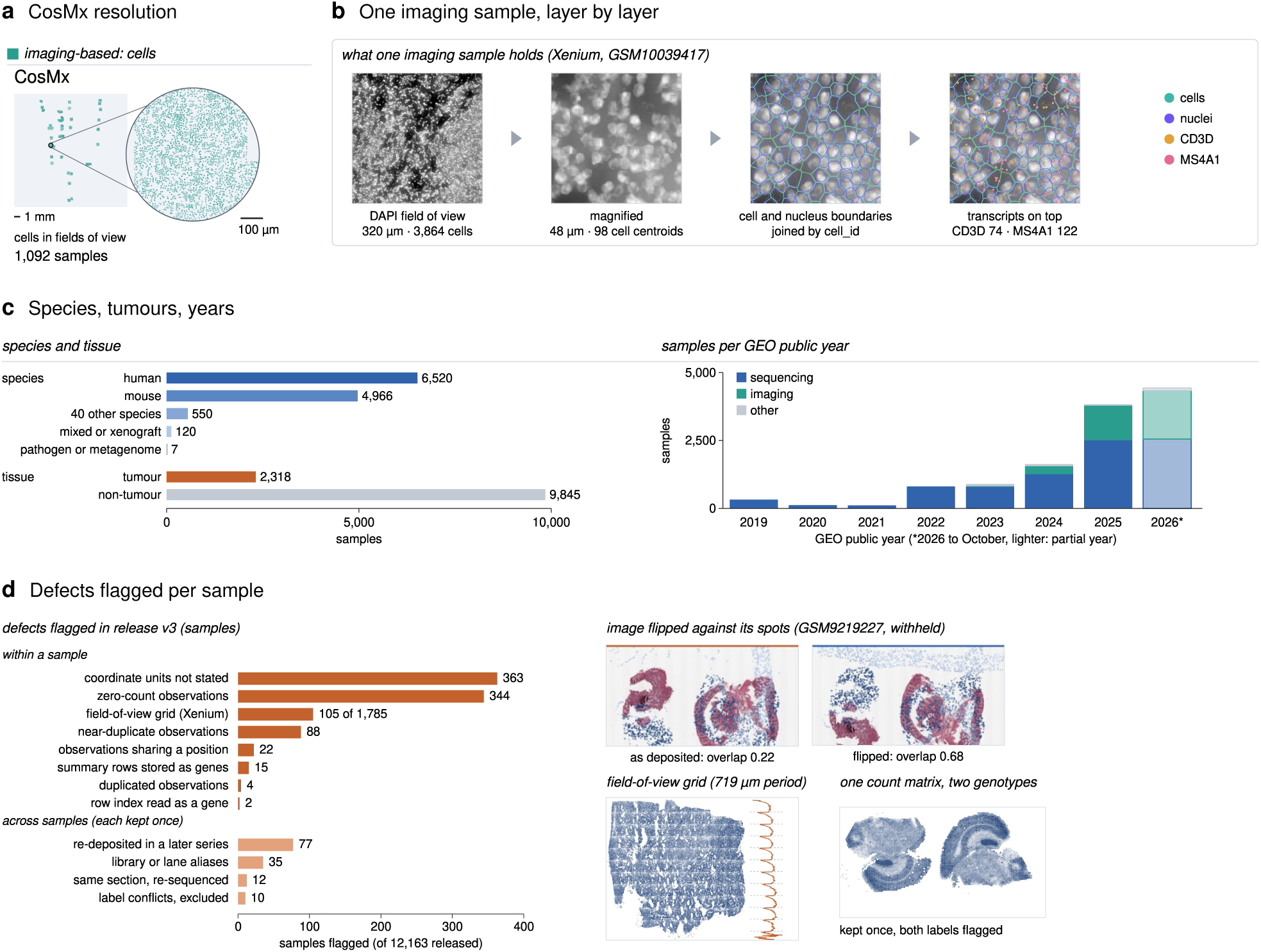
Resource details, related to Figure 3. (A) CosMx resolution. (B) Layers of one imaging sample (Xenium GSM10039417): cells (teal), nuclei (violet), and CD3D and MS4A1 transcripts. (C) Species, tumour status and samples by GEO public year (2026 shown to October in a lighter colour). (D) Defects flagged in the release: samples flagged for each kind of defect within a sample and across samples (each counted once), with three real cases.

**Figure S3.**
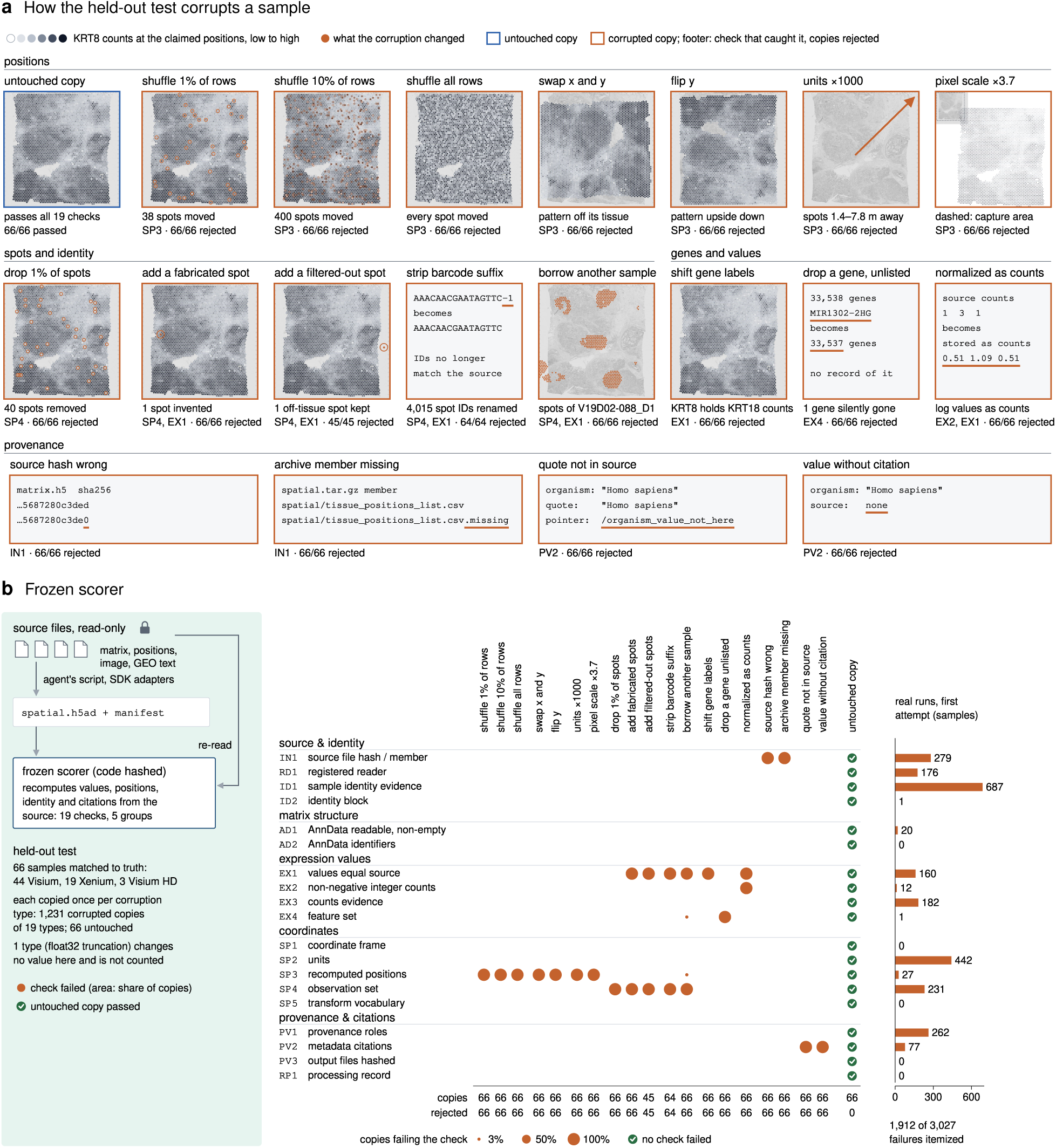
How the gate is tested, related to Figure 2. (A) Every corruption of the held-out test applied to one real sample (10x Visium breast cancer section V1_Breast_Cancer_Block_A_Section_2) with the test’s own code and random seed. Tiles show KRT8 counts at the claimed positions on the sample’s own H&E image (grey to black, low to high), and orange marks what the corruption changed. Below each tile are the check that caught it and the number of corrupted copies rejected across all 66 held-out samples. Corruptions that do not apply to a sample were not counted (21 for the filtered-out spot, 2 for the barcode suffix). (B) The frozen scorer on the held-out test set: corruption types (columns) against checks (rows), with first-attempt failures per check in real runs.

**Figure S4.**
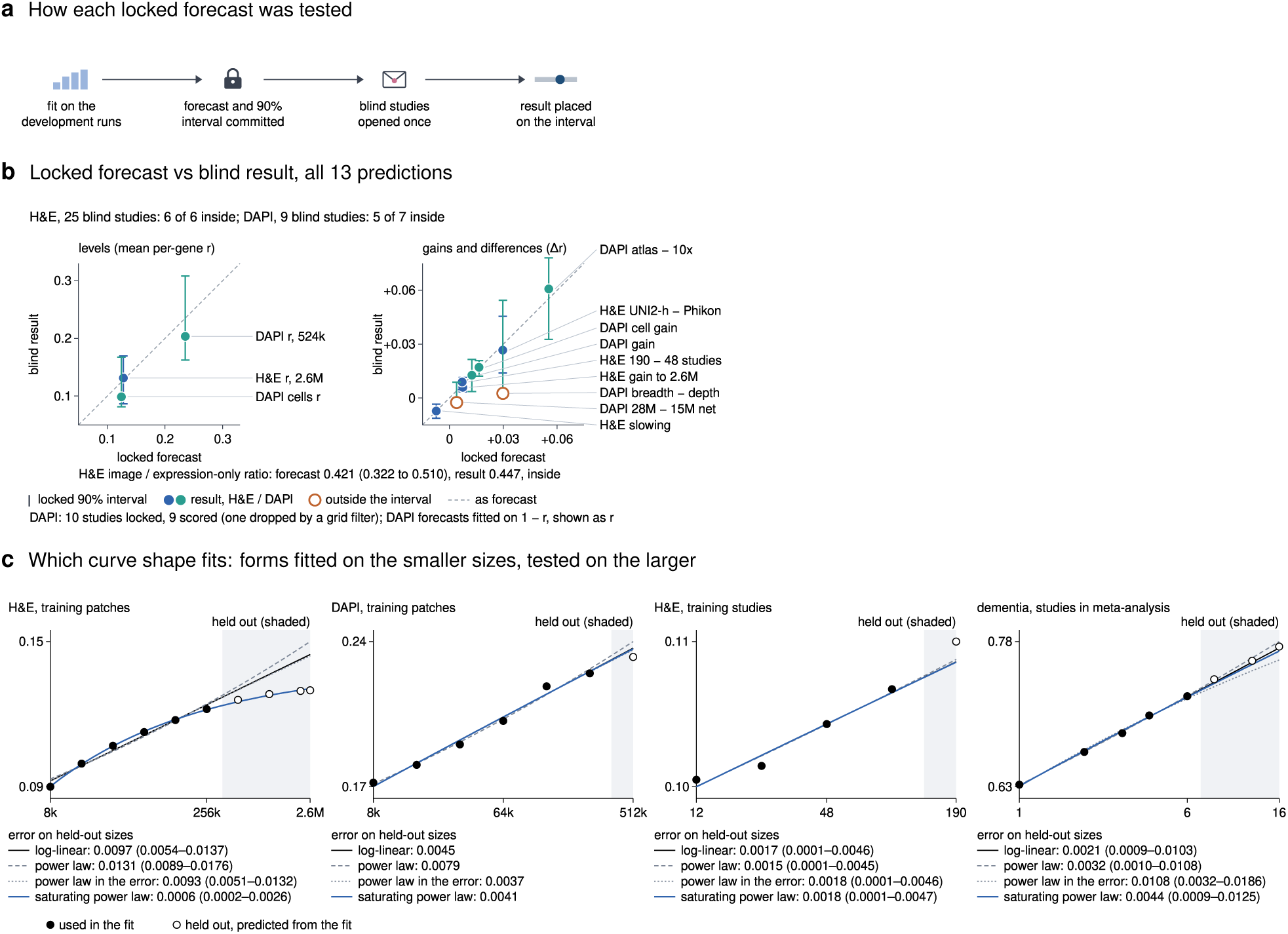
All 13 locked forecasts and the shape of the curves, related to Figure 4. (A) Test setup of each forecast. (B) Locked forecasts against results on blind studies (vertical bars, locked 90% intervals). For H&E, 6 of 6 results fell inside the interval; for DAPI, 5 of 7 did, with the breadth and network-size contrasts below the interval. (C) Which curve shape fits. Log-linear, power-law, power-law-in-the-error and saturating power-law curves were fitted on the smaller sizes of each curve (filled dots) and used to predict the larger sizes (open dots, shaded). Below each plot is the mean absolute error on the held-out sizes (95% CI over test studies; DAPI has study means only). H&E patches: UNI2-h, 22 new studies, fitted on 8k–262k. DAPI patches: fitted on 8k–262k, with 524k measured later. H&E studies: 12 to 190 studies at 65,536 patches, fitted on 12–96. Dementia: direction agreement in 28 held-out studies, fitted on 1–6 studies. Only the H&E patch curve separates the forms: the saturating power law (ceiling 0.136, 95% CI 0.105–0.174) predicts its larger sizes.

**Figure S5.**
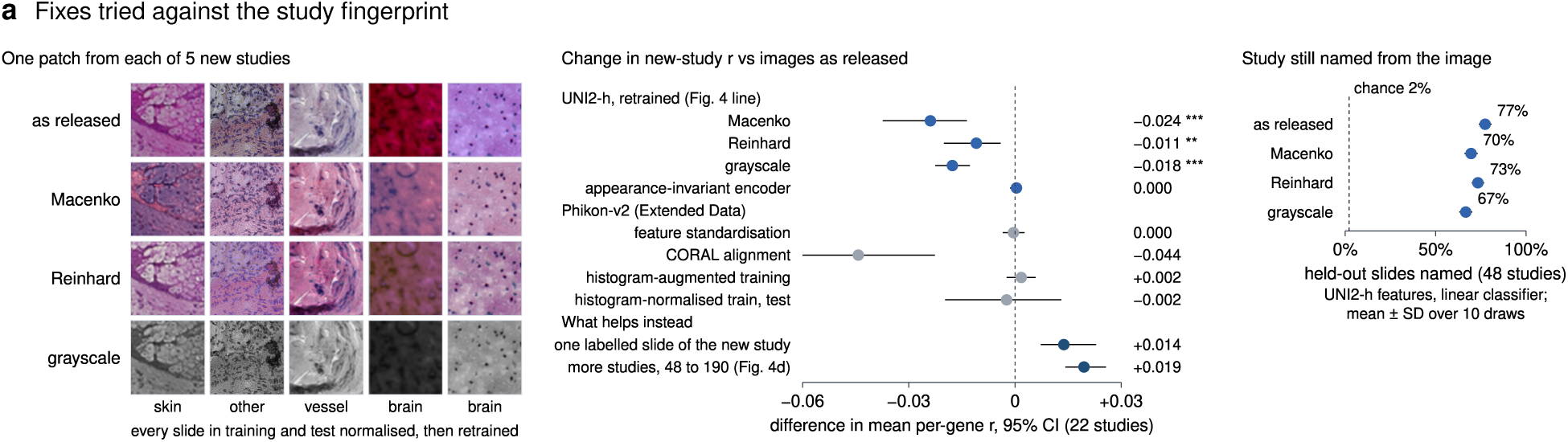
Every fix tried against the study fingerprint, related to Figure 5. Change in new-study *r* for each method tried: stain normalization, an appearance-invariant encoder, feature standardization, CORAL alignment, and histogram augmentation and normalization. These are compared with adding one labelled slide from the new study and with adding more training studies. Also shown is how often the source study can still be identified from the features.

**Figure S6.**
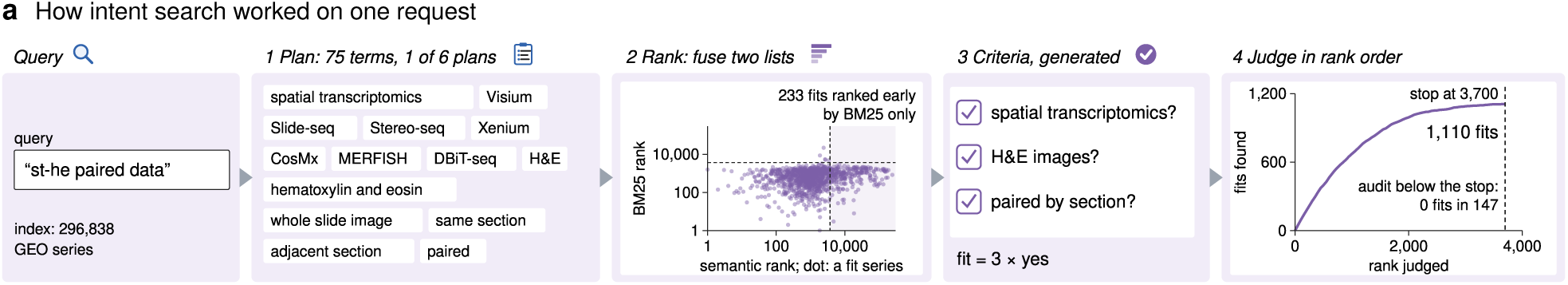
Intent-based search mechanics, related to Figure 6. An example run for “st-he paired data”: search plan terms, semantic and BM25 rankings of every candidate, generated criteria, and the judged curve. The search used an adaptive stopping rule and a tail audit, which found 0 fits in 147 audited series below the stopping point.

**Figure S7.**
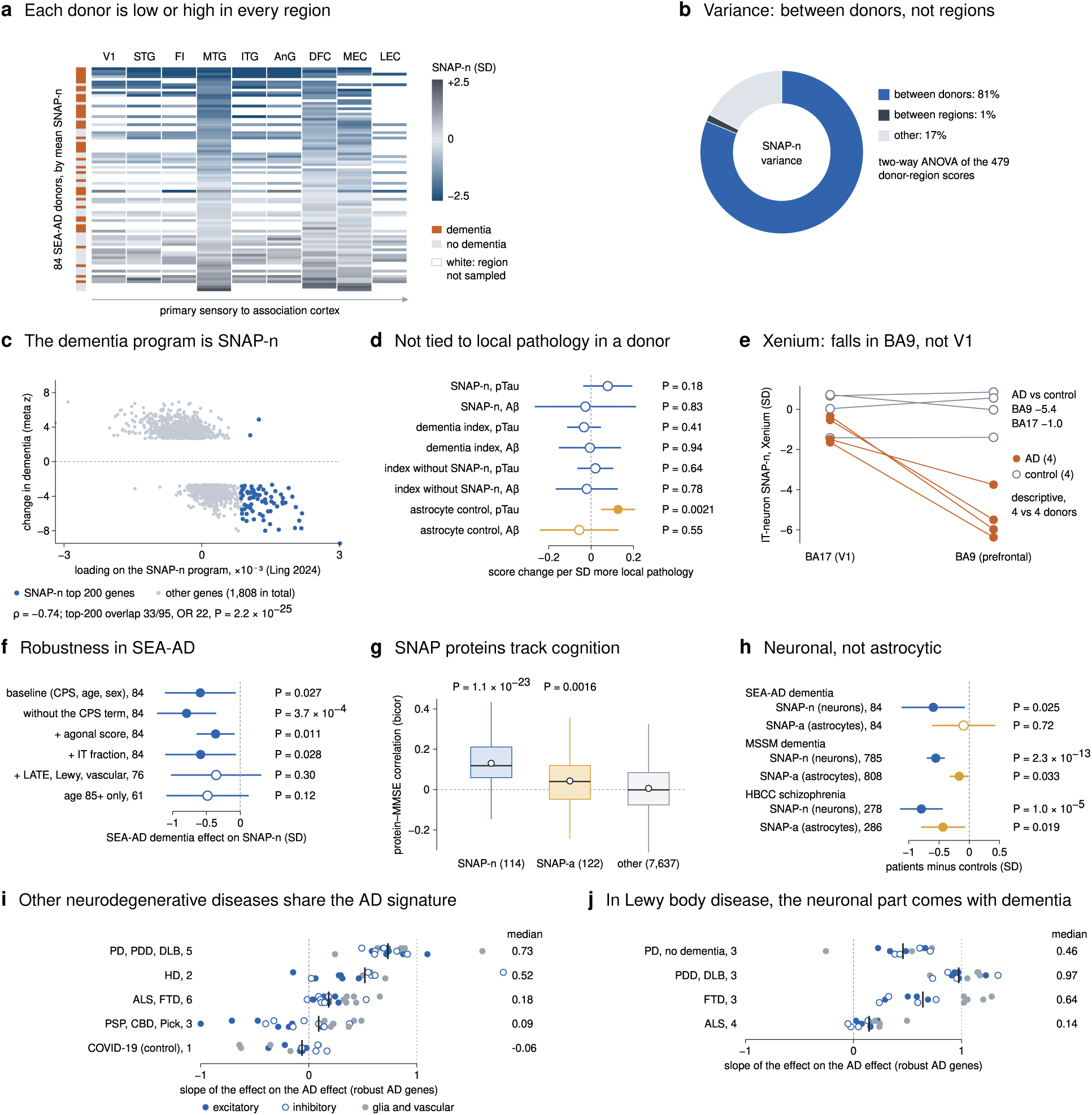
SNAP-n in detail, and how much of the AD signature other neurodegenerative diseases share, related to Figure 6. (A,B) SNAP-n scores per SEA-AD donor and brain region; 81% of the variance lies between donors. (C) Dementia programme against SNAP-n loadings. (D) No within-donor link to local pathology, whereas an astrocyte control tracks local pTau. (E) Xenium in BA9 and V1 (4 against 4 donors, descriptive). (F) Robustness analyses in SEA-AD. (G) Correlation of SNAP proteins with cognition in an independent proteome. (H) Cell-type specificity: a neuronal rather than astrocytic signal. (I) Sharing of the AD signature with other diseases (PD/PDD/DLB 0.73, HD 0.52, ALS/FTD 0.18, PSP/CBD/Pick 0.09, COVID-19 *−*0.06). (J) In Lewy body disease, the neuronal component appears only with dementia.

**Figure S8.**
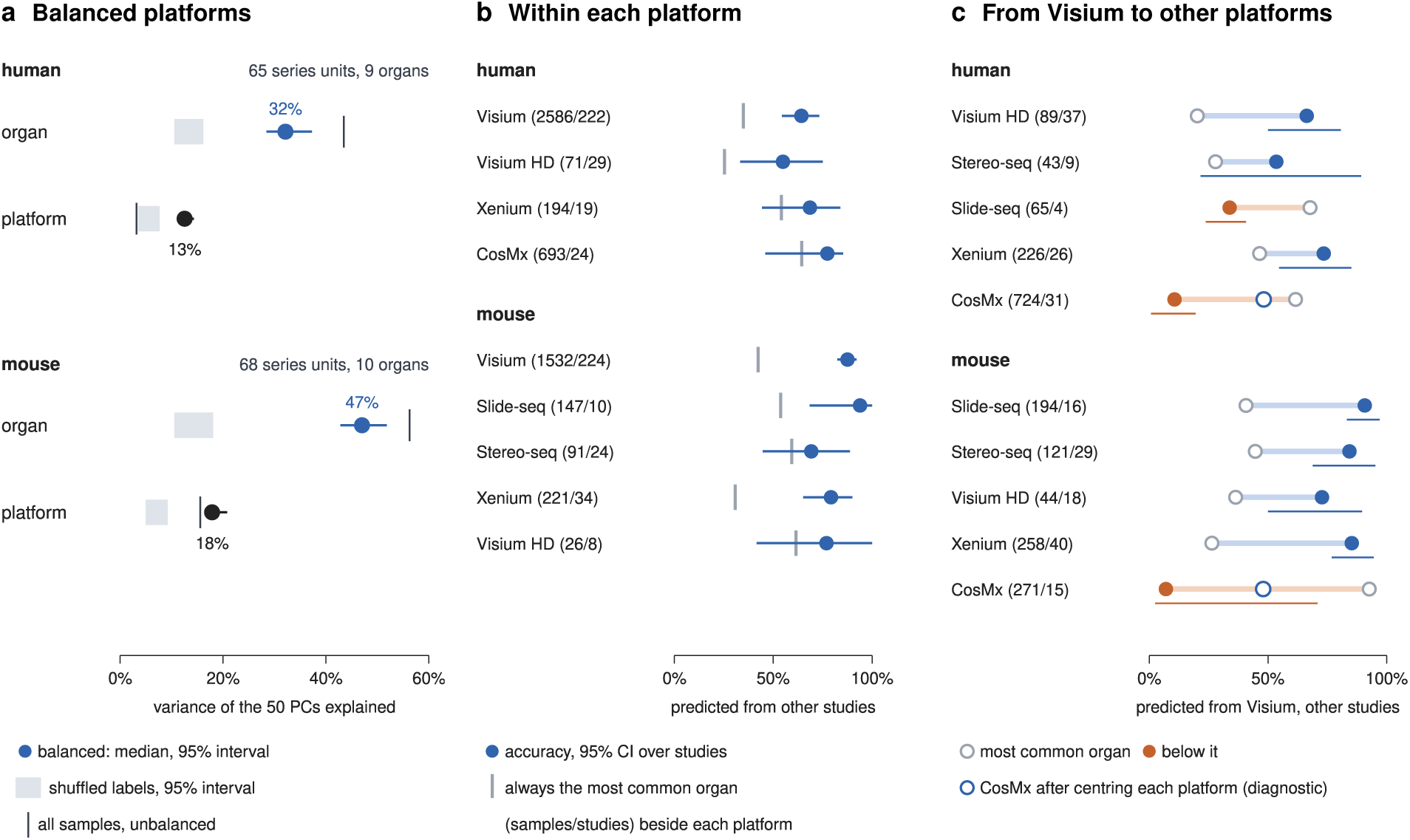
Organ against platform with platforms balanced, within platforms and from Visium to other platforms, related to Figure 3. A) Variance of the first 50 principal components of pseudobulk profiles (sequencing-based samples, no batch correction) explained by organ and by platform, with each organ contributing the same number of series-level units per platform. Organs were measured on at least two platforms, each in at least two series (human: 65 units over 9 organs; mouse: 68 over 10; PCA refitted on each of 200 draws). Dots and lines, median and 95% interval over draws; grey band, labels shuffled within the other factor; vertical ticks, all samples unbalanced (human: 92% Visium; organ 43% and platform 3%; mouse: 56% and 16%). Human: organ 32% (28–37%; chance 13%), platform 13% (11–14%; chance 5%). Mouse: organ 47% (43–52%; chance 14%), platform 18% (17–21%; chance 7%). (B) Leave-one-study-out organ prediction within each platform from the 15 nearest samples of other series (cosine distance). Prediction used genes measured in every retained sample of the platform (human Xenium: 40 genes shared by 194 of 1,074 samples; human CosMx: 770; mouse Xenium: 87). Dots, accuracy; lines, 95% CI from resampling studies; grey ticks, accuracy of always predicting the most common organ; brackets, samples/studies. Mouse CosMx (two organs, 95% liver) is omitted. Balanced accuracy: human Visium 47%, Visium HD 46%, Xenium 51%, CosMx 44%; mouse Visium 74%, Slide-seq 84%, Stereo-seq 47%, Xenium 65%, Visium HD 74%. (C) Organ predicted for samples of each platform from their 15 nearest Visium samples of other series, on genes shared with Visium. Open grey symbols, share of the most common organ among query samples; filled symbols, accuracy (orange when below the grey symbol) with 95% CI over studies. CosMx profiles lie far from Visium, at 1.6–1.8 times the spread between organ means, and are mostly predicted as a single organ. Centring each platform on its own mean (open blue, diagnostic only) raises accuracy to 48% in both species, whereas Xenium, also imaging-based, transfers (74% human, 85% mouse).

**Figure S9.**
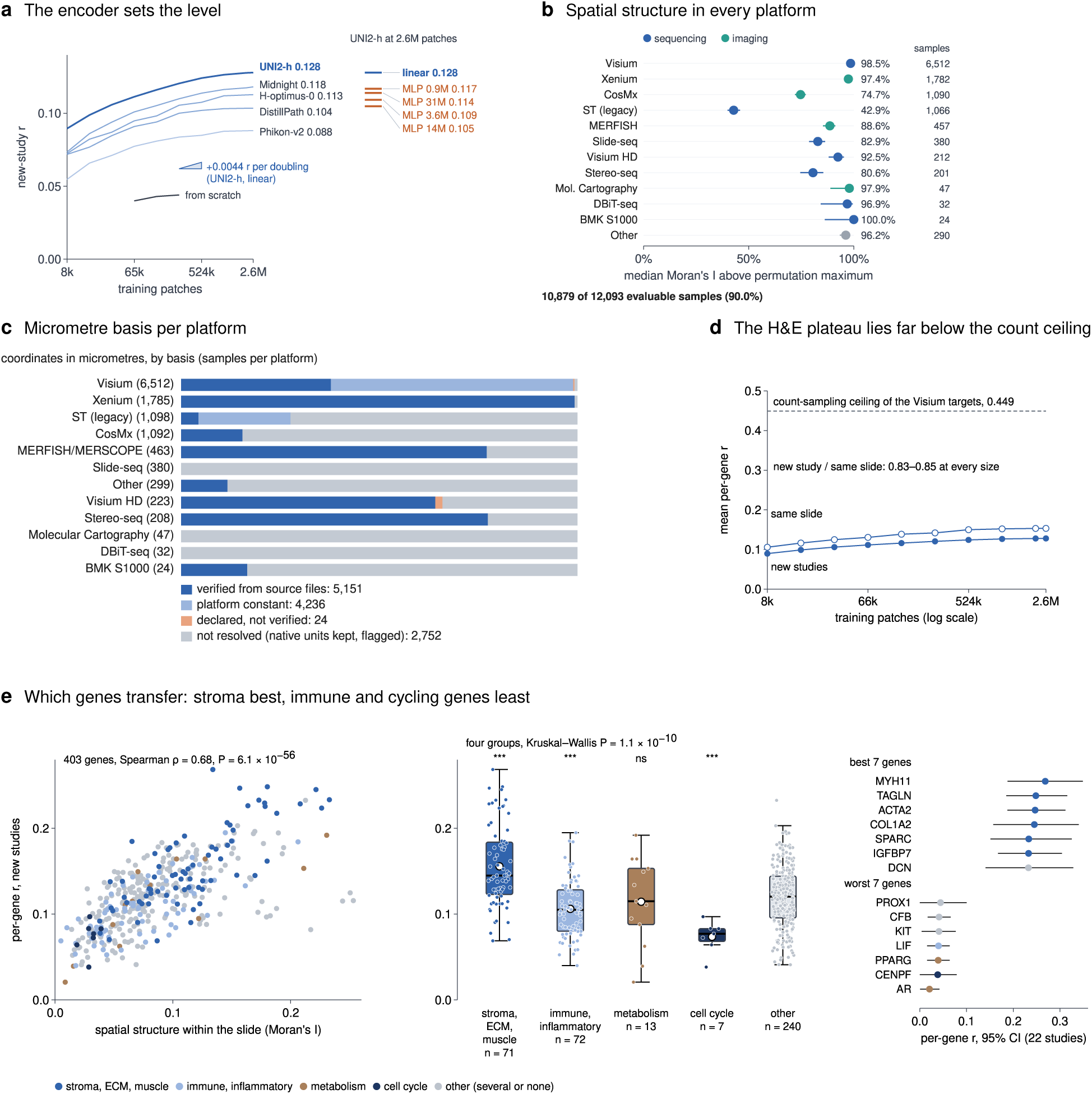
Supporting analyses, related to Figures 3 and 4. (A) Five frozen encoders with a linear read-out. Right column: read-outs on frozen UNI2-h features trained with 2.6 million patches, the linear read-out (blue) against four MLP read-out heads (orange, labelled by parameter count); none outperformed the linear head. The triangle, drawn to scale, shows the gain of the UNI2-h linear curve per doubling of training patches. (B) Spatial structure on every platform. (C) Micrometre basis by platform. (D) New-study (filled) and same-slide (open) *r* of UNI2-h against training patches, with the count-sampling ceiling of the Visium targets (dashed; Figure 5F); the ratio of the two stays at 0.83–0.85. (E) Which genes transfer: per-gene new-study *r* against spatial autocorrelation within a slide (Moran’s I [89]), by MSigDB Hallmark programme [90], with the best and worst genes.

